# Dscam homophilic specificity is generated by high order *cis*-multimers coupled with *trans* self-binding of variable Ig1 in Chelicerata

**DOI:** 10.1101/2019.12.15.877159

**Authors:** Fengyan Zhou, Guozheng Cao, Songjun Dai, Guo Li, Hao Li, Zhu Ding, Shouqing Hou, Bingbing Xu, Wendong You, Feng Shi, Xiaofeng Yang, Yongfeng Jin

**Affiliations:** MOE Laboratory of Biosystems Homeostasis & Protection and Innovation Center for Cell Signaling Network, College of Life Sciences, Zhejiang University, Hangzhou, Zhejiang, ZJ310058, China; Department of Neurosurgery, First Affiliated Hospital, School of Medicine, Zhejiang University, Hangzhou, Zhejiang, ZJ310058, China

**Keywords:** Down syndrome cell adhesion molecule, isoform diversity, alternative promoter, homophilic binding, multimer, combinatorial specificity, cell identity, self-recognition, self-avoidance, self–non-self discrimination, Chelicerata

## Abstract

By alternative splicing, *Drosophila Down syndrome cell adhesion molecule* (*Dscam1*) encodes tens of thousands of proteins required for establishing neural circuits, while Chelicerata encodes a family of ∼ 100 shortened *Dscam* (sDscam) isoforms via alternative promoters. We report that Dscam isoforms interact promiscuously *in cis* to generate a vast repertoire of combinatorial homophilic recognition specificities in Chelicerata. Specifically, sDscams formed high order *cis*-multimers without isoform specificity involving the membrane-proximal fibronectin type III (FNIII) 1-3 and transmembrane (TM) domains and associated specifically *in trans* via antiparallel self-binding of the first variable immunoglobulin (Ig1) domain. We propose that such sDscam combinatorial homophilic specificity is sufficient to provide each neuron with a unique identity for self–non-self discrimination. In many respects, our results amazingly mirror those reported for the structurally unrelated vertebrate protocadherins (Pcdh) rather than for the closely related fly Dscam1. Thus, our findings blur the distinction between the neuronal self-avoidance of invertebrates and vertebrates and provide insight into the basic principles and evolution of metazoan self-avoidance and self–non-self discrimination.

## Introduction

Neuronal self-avoidance refers to the tendency of neurites from the same neuron to avoid each other, which is conserved in vertebrates and invertebrates and plays a vital role in neural circuit assembly ^1, 2^. Also, this process is mediated by strictly homophilic interactions between cell recognition proteins that trigger a neurite-repulsive rather than -adhesive signal. In turn, neuronal self-avoidance requires a molecular mechanism by which each cell discriminates self from non-self ^1, 2^. Thus, one challenge in deciphering the basis of self-avoidance is elucidating how neurons discriminate self from non-self. One solution to this challenge is to endow each neuron with a unique identity by expressing diverse families of cell adhesion molecules ^1,2,3^.

In *Drosophila*, neuronal self-avoidance is mediated by extraordinary cell recognition molecules of the immunoglobulin superfamily, which are encoded by the *Down syndrome cell adhesion molecule* (*Dscam1*) locus via alternative splicing ^1, 3^. This vast repertoire of Dscam1 recognition molecules is sufficient to bestow a unique molecular identity on each neuron and homophilic interaction specificity, thereby allowing neuronal processes to distinguish between self and non-self ^4, 5^. If neurites of the same neuron approach each other, neurites with identical Dscam1 isoforms exhibit homophilic interactions, resulting in self-avoidance due to contact-dependent repulsion. By contrast, neurites from different neurons express distinct Dscam1 protein repertoires that do not engage in homophilic binding and thus contact each other ^1, 6^.

Stochastic alternative splicing of *Drosophila Dscam1* enables it to encode up to 38,016 distinct isoforms, each of which comprises 1 of 19,008 distinct ectodomains linked to one of two alternative transmembrane regions ^7,8,9,10^. Individual neurons stochastically express a unique set of distinct Dscam1 isoforms, which may engage in highly isoform-specific homophilic binding between neurons, thus endowing each neuron with a unique molecular identity ^4, 5^. Dscam isoform diversity plays essential roles in self-recognition and self-avoidance ^11,12,13^. In contrast to insect *Dscam1*, vertebrate *Dscam* genes do not produce extensive protein isoforms, and functional studies revealed that mouse *Dscam* genes are not essential for neuronal self-avoidance ^14, 15^.

However, another set of genes, the clustered *Pcdh*s, perform a similar function in vertebrates, and generate enormous cell-surface structural diversity ^1,2,3, 16^. Pcdhs belong to the cadherin superfamily, the largest and best-established family of cell-adhesion molecules. In human and mouse, 53 and 58 Pcdh proteins are encoded by three tandemly arranged gene clusters of *Pcdhα*, *Pcdhβ*, and *Pcdhγ*, respectively ^17, 18^. In contrast to fly Dscam1, the individual variable exon is preceded by an alternative promoter, and differential expression of Pcdh isoforms is achieved by a combination of stochastic promoter selection and alternative splicing ^19,20,21^. Like fly Dscam1 isoforms, almost all clustered Pcdh proteins engage in isoform-specific *trans* homophilic interactions ^22, 23^. However, in contrast to fly Dscam1 isoforms, which act as a monomer, Pcdh proteins act as multimeric recognition units to expand the adhesive interface ^22,23,24,25,26,27,28,29^, which provides a reasonable explanation for the different isoform number between vertebrate Pcdh and fly Dscam1. Deletion of the Pcdh γ-subcluster or all three clusters caused self-avoidance defects of dendrites and axons ^30, 31^, positing that *Drosophila* Dscam1 isoforms and vertebrate clustered Pcdhs employ similar strategies for self-avoidance. Also, vertebrate and *Drosophila* neurons have solved the self-avoidance problem based on a fundamentally similar principle albeit with a different set of recognition molecules.

Of particular relevance to the remarkable functional convergence of *Drosophila* Dscam1 and vertebrate clustered Pcdhs is our recent discovery of a “hybrid” gene family in the subphylum Chelicerata. This gene family was composed of shortened *Dscam* genes with tandemly arrayed 5’ cassettes, which encoded ∼ 50–100 isoforms varying across various Chelicerata species via alternative promoter ^32, 33^. Since these Chelicerata Dscams lack the N-terminal Ig1–6, 10 domains and FNIII3–4, 6 domains present in classical DSCAM, we refer to this type of Dscam as shortened *Dscam* (sDscam) to distinguish it from classical Dscam. Based on their different variable 5’ cassettes encoding one or two Ig domains, these sDscams can be subdivided into the sDscamα and sDscamβ subfamilies. Thus, all sDscam isoforms share the same domain structure but contain variable amino acid sequences within the N-terminal and two Ig domains in the extracellular region. Interestingly, the 5’ variable region of Chelicerata *sDscams* shows remarkable organizational resemblance to that of vertebrate-clustered Pcdhs ^14, 32, 33^. Similar to *Drosophila Dscam1* and vertebrate Pcdhs, Chelicerata *sDscam* are abundantly expressed in the nervous system ^32, 33^. Because Chelicerata sDscams are remarkably similar to Drosophila Dscam1, and exhibit a remarkable organizational resemblance to the vertebrate-clustered Pcdhs, with the latter two proteins both capable of mediating self-recognition and self-avoidance, we speculate that these sDscam isoforms play analogous roles in Chelicerata species. Therefore, a systemic examination of the homophilic recognition specificities of these clustered sDscam isoforms was performed to address their roles in specifying single-cell identity and neural circuit assembly.

In this study, we show that most clustered sDscams can engage in highly specific homophilic interactions via antiparallel self-binding of the variable Ig1 domain. Moreover, we provide compelling evidence that sDscam isoforms can associate promiscuously as high order *cis*-multimers, which is mediated by the constant FNIII1–3 and transmembrane domains. Based on a large body of experimental evidence and structural modeling, we concluded that sDscams mediate self-recognition via promiscuous *cis* interactions coupled with strictly homophilic Ig1/Ig1 interactions in *trans*, possibly forming an interconnected latticed protein assembly between apposed cell surfaces. We propose that these sDscam homophilic specificities are sufficient to provide the unique single-cell identity necessary for neuronal self–non-self discrimination. Interestingly, in many respects, Chelicerata sDscams show more remarkable parallels with the genetically unrelated vertebrate Pcdhs than to the closely related fly Dscam1. Thus, our findings provide mechanistic and evolutionary insight into self–non-self discrimination in metazoans and enhance our understanding of the general biological principles required for endowing cells with distinct molecular identities.

## Results

### Cluster-wide analysis of sDscam-mediated homophilic interactions

The *Mesobuthus martensii sDscam* gene clusters encode diverse cell-adhesion proteins: 40 alternate sDscamα isoforms and 55 sDscamβ1, β2, β3, β4, β5, and β6 isoforms (Fig. 1a) ^32, 33^. To investigate whether sDscam isoforms mediate homophilic binding, we expressed the sDscam proteins in Sf9 cells using an insect baculovirus expression system (Fig. 1b). This system is a powerful tool for investigating homophilic interactions between cell surface adhesion molecules ^34^. We prepared Sf9 cells expressing sDscam, as well as Sf9 cells infected with the parental virus as a negative control and expressing fly Dscam1 as a positive control. Sf9 cells expressing constructs encoding full-length sDscamβ6v2 (β6v2FL-mCherry) or sDscamβ6v2 lacking the cytoplasmic domain (β6v2Δcyto-mCherry) exhibited strong cell aggregation (Supplementary Fig. 1a), indicating that the homophilic interaction was mediated by sDscamβ6v2 *in trans* independently of the cytoplasmic region. Because our results indicated that deletion of the cytoplasmic tail of sDscam did not significantly affect the formation of cell aggregates (Supplementary Fig. 1a), we used Δcyto constructs for all sDscam proteins in the cell aggregation assay.

**Fig. 1.**
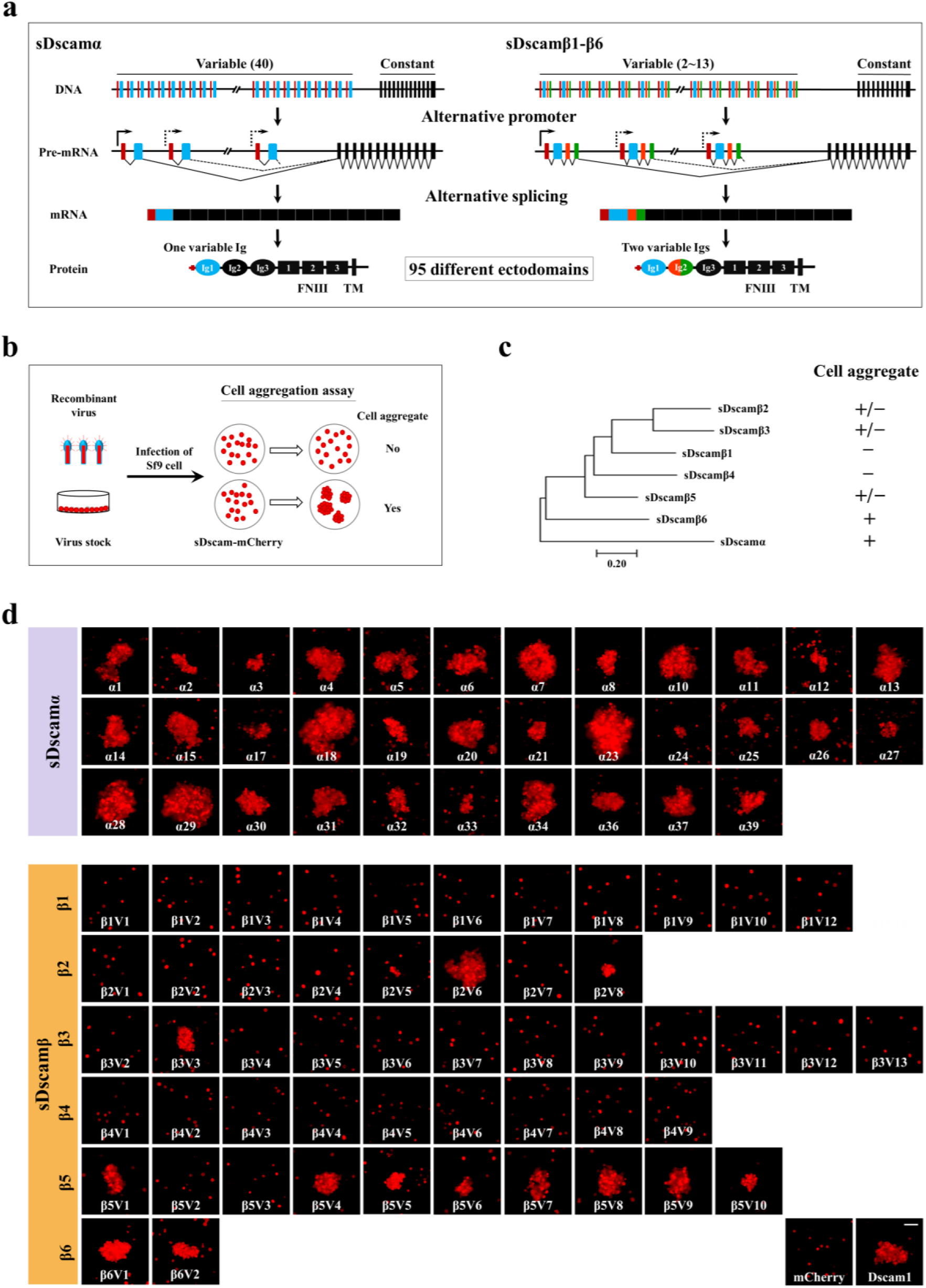
Cluster-wide analysis of sdscam-mediated homophilic binding in *M. martensii*. (a) Overview of the *M. martensii* sDscam gene clusters. Variable exons (coloured) are joined via *cis*-splicing to the constant exons (black) in sDscamα (left) and sDscamβ1–β6 (right) encodes Ig1–2 domains. The constant exons of sDscamα and sDscamβ encode the Ig2–3 or Ig3 domains, FNIII1–3 domains, the transmembrane and cytoplasmic domains. (b) Schematic diagram of the cell aggregation assay. mCherry-tagged sDscam proteins were expressed in Sf9 cells for assaying their ability to form cell aggregates. As shown in the diagram, cells expressing some sDscam-mCherry alone did not aggregate as negative control - mCherry, while strong cell aggregates were observed with cells expressing other sDscam-mCherry as positive control Dscam1-mCherry. (c) The summary of results for homophilic binding properties. An evolutionary relationship among distinct sDscam subfamilies is shown on the left. (d) The outcome of cell aggregation of 85 sDscam isoforms when assaying individually. mCherry and fly Dscam1 isoform were expressed as negative and positive control. See also Supplementary Fig. 1b. Scale bar, 100 µm.

Except for a few sDscam cDNAs that failed to be cloned possibly due to low expression, we performed a systematic analysis of the homophilic interactions of 85 of the 95 sDscam proteins (sDscamα, sDscamβ1–β6). We found that the feature or size of the cell aggregates varied markedly across sDscam subfamilies (Fig. 1c, d, Supplementary Fig. 1b). Cells expressing each alternate sDscamα showed extensive aggregation for all isoforms tested (Fig. 1d, lane 1-3). By contrast, cells expressing most of the alternate sDscamβ1–β4 isoforms failed to form aggregates, except for a few isoforms (Fig. 1d, lane 4-7). In particular, none of the alternate sDscamβ1 or β4 isoforms formed aggregates. Moreover, the aggregates of cells expressing sDscamβ isoforms were smaller than those expressing sDscamαs (Fig. 1d, Supplementary Fig. 1b). These observations suggest that the ability of each sDscam isoform to mediate homophilic aggregation differs in a cluster-specific manner.

Notably, cells expressing individual sDscam isoforms from the same cluster, which differed only in the N-terminal variable region, exhibited markedly different cell aggregation activity. For example, sDscamβ2v6 and β2v8 formed homophilic aggregates, but other members of the sDscamβ2 subfamily did not (Fig. 1d, lane 5). Similar results were obtained when assaying individually sDscamβ3 and β5 isoforms (Fig. 1d, lane 6, 8). This discrepancy in the aggregation activity between isoforms from the same or different clusters was likely the consequence of differences in the expression, membrane localization, or intrinsic *trans*-binding affinities of individual sDscam isoforms ^23^. Failure to form cell aggregates in mammalian Pcdhα isoforms is reportedly due to the lack of membrane localization ^23, 35, 36^. However, this can be ruled out because immunostaining indicated that sDscamβ4v3 was present on the surface of Sf9 cells, as was sDscamα14 (Supplementary Fig. 1c, panels i and iii). Therefore, we speculate that the different aggregation activities likely reflect, at least in part, differences in the intrinsic *trans* homophilic binding affinities of the individual isoforms.

### The first two N-terminal Ig domains are required for homophilic *trans* binding

To explore how sDscam proteins mediated differential homophilic *trans-*binding, we firstly defined the minimum domain required for homophilic interactions. To this end, a series of N-terminal truncations of the extracellular domain of *sDscamα*14 fused with mCherry were subjected to cell aggregation assay (Fig. 2a). We did not observe cell aggregation for each of these deletion constructs in which the first one to five domains were successively deleted from sDscamα14 (Fig. 2a, panels ii-vi). Similarly, all constructs lacking the Ig1 domain did not show aggregation for sDscamβ3v3 and β6v1 (Fig. 2a, panels viii-xii, Supplementary Fig. 2a, panels 2-6). Of all the truncated mutants in three independent assays, homophilic binding activity was dependent on the presence of the Ig1 domain (Fig. 2b). These results indicate that the first N-terminal domain (Ig1) is required for sDscam-mediated homophilic *trans* binding.

**Fig. 2.**
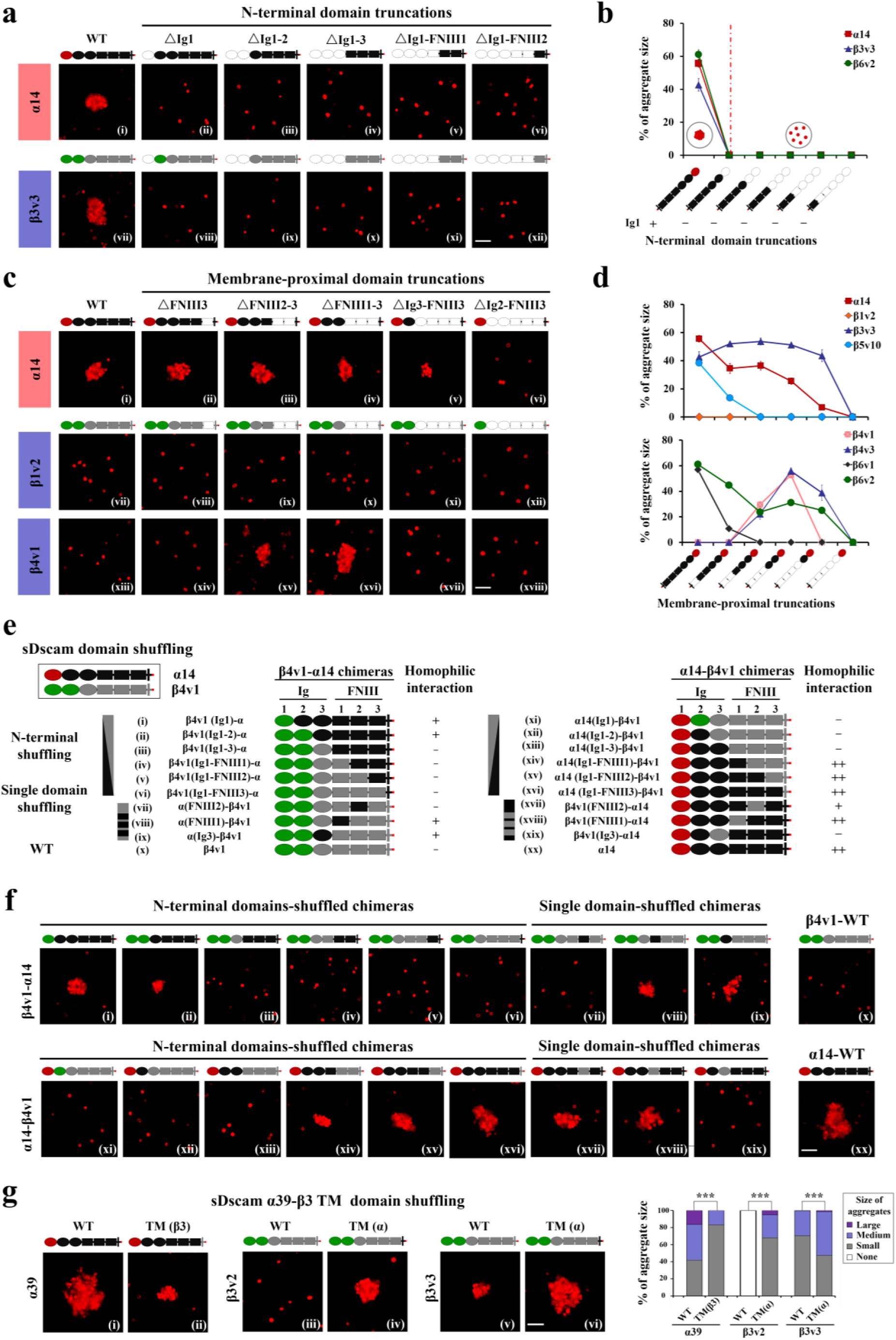
Homophilic *trans*-binding is associated with variable and constant domains of sDscam. (a) A series of N-terminal truncations of the extracellular domain of *sDscam* fused with mCherry were examined for cell aggregation assay. Three independent assays were performed in sDscamα14, sDscamβ3v3 and sDscamβ6v2, respectively. All of the sDscam truncations lacking the N-terminal Ig1 domain failed to form cell aggregate. See also Supplementary Fig. 2a. (b) The ratio change of cell aggregate size (medium and large) of sDscam N-terminal truncations. Results were obtained from three independent experiments and expressed as mean ± SEM. (c) The first two N-terminal domains are required for *trans* homophilic binding. Eight sDscam isoforms including 1 sDscamα and 7 sDscamβs isoforms were deleted starting with the membrane-proximal FNIII3 domain, and these truncations were tested for cell aggregation. These data showed that all of the truncated constructs containing only one Ig1 domain failed to form cell aggregates, and the smallest truncations that exhibited binding ability contain N-terminal Ig1–2 domains. Supplementary Fig. 2a. (d) Homophilic *trans*-binding is associated with constant extracellular domains of sDscam. The graph showed that cell aggregation size (medium and large) of many sDscam truncations changed obviously, except sDscamβ1v1. See also Supplementary Fig. 2a. Results were obtained from three independent experiments and expressed as mean ± SEM. (e) Schematic diagrams for domain shuffling mutants used in the experiments, along with a summary of outcomes from homophilic interaction assays. Extracellular domain of sDscamα14 (red and black) were replaced with the corresponding domains of sDscamβ4v1 (green and gray), or vice versa. All of the chimeras bearing the Ig3–FNIII1 domain from sDscamα14 (black) could affect cell aggregation. See also Supplementary Fig. 2b, c. (f) Cell adhesion images were presented corresponding to the shuffled chimeras shown in Fig. 2e (i–xx). These results indicate that homophilic binding is regulated by constant extracellular domains of sDscam. (g) The TM domain is associated with homophilic binding. TM domain shuffling experiments proportion of each category of the quantification of the cell aggregation corresponding to the shuffled chimeras (panels i–vi). Results were obtained from three independent experiments and expressed as mean. Mann-Whitney U-test was performed to determine significance. ***p < 0.001. Scale bar, 100 µm.

We produced a series of truncations in which extracellular domains were successively deleted starting with the membrane-proximal FNIII3 domain (Fig. 2c). These truncations harboured one sDscamα and seven sDscamβ isoforms, four of which were derived from two clusters, enabling comparison of various isoforms from the same cluster or from different clusters (Fig. 2c, Supplementary Fig. 2a). Representative examples of truncation are shown for sDscamα, sDscamβ1v2, and β4v1 in Fig. 2c. Five additional examples for sDscamβ3v3, β4v3, β5v10, β6v1, and β6v2 are shown in Supplementary Fig. 2a. With the exception of the sDscamβ1v2 truncated constructs, all of which did not induce cell aggregation as the complete ectodomains, at least one truncated construct containing the first two N-terminal Ig domains mediated cell aggregation for each sDscam tested (Fig. 2c, panels i–xviii, Supplementary Fig. 2a, panels 7–36), while all truncated constructs containing only the Ig1 domain did not (Fig. 2c, panels vi, xii and xviii, Supplementary Fig. 2a, panels 12, 18, 24, 30 and 36). These results indicate that the smallest protein that exhibited *trans* interaction contains two N-terminal Ig1– 2 domains (Fig. 2c, d, Supplementary Fig. 2a), compatible with the fact that the variable Ig1 domain (possibly in combination with Ig2) acted as a *trans*-binding interface (see results below). Thus, the first two N-terminal Ig domains are essential for sDscam-mediated homophilic *trans* binding.

### Homophilic *trans*-binding is associated with the constant extracellular and transmembrane domains of sDscam

Based on data from eight independent assays with successive truncation from the membrane-proximal extracellular domain (Fig. 2c, Supplementary Fig. 2a), we observed three different changes in aggregation. First, cell aggregation was gradually decreased as the number of domains deleted increased in most sDscam isoforms tested, such as sDscam*α*14, sDscamβ5v10, β6v1, and β6v2 (Fig. 2c, panels i–vi, Supplementary Fig. 2a, panels 19-36). In these cases, constant domains might enhance sDscam-mediated homophilic *trans* binding. Second, deletion of two or three membrane-proximal FNIII domains in sDscamβ4v1 and β4v3 rescued their cell aggregation activity, more efficiently in β4v1ΔFNIII1–3 and β4v3ΔFNIII1–3 (Fig. 2c, panels xiii–xviii, Supplementary Fig. 2a, panels 13–18). These observations suggest that the FNIII domains of sDscamβ4 inhibit homophilic *trans* binding. Third, we found that sDscamβ1 homophilic interactions could not be rescued by deletion of any domain (Fig. 2c, panels vii– xii). Therefore, the constant domains may be associated with homophilic binding in a cluster-specific manner.

Interestingly, the change in cell aggregation varies markedly upon domain truncation even between isoforms from the same cluster. For example, we observed cell aggregates in β6v2ΔFNIII2–3, β6v2ΔFNIII1–3, and β6v2ΔIg3–FNIII3, but not in their β6v1 counterparts (Supplementary Fig. 2a, panels 27–29 and 33–35). Similarly, β4v3ΔIg3–FNIII3, but not β4v1ΔIg3–FNIII3, exhibited aggregation activity (Fig. 2c, panel xvii, Supplementary Fig. 2a, panel 17). This suggests that the constant domains influence homophilic *trans* binding by coupling with variable domains.

To further investigate how the constant extracellular domains contribute to homophilic *trans* binding, we performed experiments in which domains were shuffled between sDscamα14, which binds, and sDscamβ4v1, which does not (Fig. 2e). Constructs in which the extracellular domain of sDscamα14 was replaced by the corresponding domain of sDscamβ4v1, or *vice versa*, were produced and tested for cell aggregation activity (Fig. 2e, f). As a result, replacing the Ig2–FNIII3 or Ig3–FNIII3 constant domains of sDscamβ4v1 with the corresponding region of sDscamα14 caused homophilic binding, while chimeric constructs containing the FNIII1–3 or less-constant domains of sDscamα14 did not show homophilic binding (Fig. 2e, f, panels i–vi). By contrast, replacing the FNIII1–3 or more-constant domains of sDscamα14 by the corresponding region of sDscamβ4v1 failed to induce cell aggregation, while chimeric constructs containing the FNIII2–3 or FNIII3 domain of sDscamβ4v1 resulted in cell aggregation (Fig. 2e, f, panels xi–xvi). The fact that both truncated and shuffled constructs derived from sDscamβ4v1 could mediate homophilic interactions indicates that there is no inherent physical barrier, such as binding incompatibility at the *trans* interface, preventing homophilic recognition of these proteins (Fig. 2c, panels xv and xvi, Fig. 2f, panel i and ii). Taken together, these data demonstrate that, in addition to matching between the N-terminal variable domains, the constant domains may affect homophilic *trans* binding.

To further determine which extracellular domain contributed to homophilic *trans* binding, we carried out chimeric mutation experiments in which a single domain was shuffled between sDscamα14 and sDscamβ4v1. Replacement of the Ig3 or FNIII1 domain of sDscamβ4v1 by the corresponding region of sDscamα14 caused detectable homophilic binding (Fig. 2f, panels viii and ix). Conversely, individual replacement of the Ig3–FNIII2 of sDscamα14 by the corresponding domain of sDscamβ4v1 decreased or even abolished homophilic binding activity (Fig. 2f, panels xvii–xix). Therefore, the Ig3 and FNIII1 domains play an important role in homophilic *trans* binding. Similar domain shuffling experiments were performed between sDscamα14 and sDscamβ1v1 (Supplementary Fig. 2b). However, individual or combined replacement of the constant extracellular domain of sDscamβ1v1 by the corresponding region of sDscamα14 did not cause detectable aggregation (Supplementary Fig. 2b, panels 3-7). Taken together, these results show that the constant extracellular domains contributed to regulating homophilic *trans* binding in a cluster-specific manner.

In addition, we performed domain-shuffling experiments between another sDscam pair: sDscamα39 (very strong homophilic binding) and sDscamβ3v2/v3 (no or weak homophilic binding) (Fig. 2g, Supplementary Fig. 2c). These shuffling experiments not only identify the extracellular domain that contributes to the homophilic *trans* binding activity of a chimera but also indicate that the TM region is involved in homophilic binding (Fig. 2g, Supplementary Fig. 2c). Strikingly, replacement of the TM domain of sDscamα39 by that of sDscamβ3 significantly decreased homophilic binding activity (Fig. 2g, panels i–ii). Conversely, replacement of the transmembrane domain of sDscamβ3v2 with the corresponding region of sDscamα39 caused detectable aggregation (Fig. 2g, panels iii–iv). Likewise, replacement of sDscamβ3v3 with the transmembrane domain of sDscamα39 significantly increased aggregation (Fig. 2g, panels v– vi). However, replacement of sDscamβ4v1 by the transmembrane domain of sDscamα14 did not affect its aggregation, and *vice versa* (Fig. 2e, f, panels vi versus x, and xvi versus xx). These results indicate that the transmembrane domain contributes to homophilic *trans* binding, at least in some sDscams.

Overall, the domain-swapping results, together with the domain-truncation experiments (Fig. 2a, c, Supplementary Fig. 2a), suggest that the extracellular and transmembrane domains might be involved in homophilic *trans* binding, with the relative contributions of each domain varying among the sDscam clusters.

### sDscams exhibit highly isoform-specific binding

To analyze the specificity of interaction between different sDscams as well as isoforms differing in the 5’ variable Ig domains, we assessed cell aggregates formed by mixing two fluorescently labeled cell populations (Fig. 3a). Each sDscam was expressed with mCherry or enhanced green fluorescent protein (EGFP) fused to the C-terminus and assayed for binding specificity. We first investigated the specificity of the interaction between sDscamα isoforms, which differ in the Ig1 domain at the N-terminus. To determine the stringency of recognition specificity, we generated pairwise sequence identity heat maps of the variable Ig1 domains (Fig. 3b). Using these heat maps, we identified sDscam pairs with the highest pairwise sequence identity within their Ig1 domains. Fourteen of the closely related sDscams (>87% identity, Fig. 3b) were tested together with 21 more distantly related sDscams (Fig. 3c). In total, we tested 35 unique pairs of sDscams with sequence identity for non-self pairs ranging from 50–97% in the Ig1 domains. For most sDscamα pairs, only self-pairs on the matrix diagonals exhibited intermixing of red and green cells, while all non-self pairs formed separate, noninteracting homophilic cell aggregates (Fig. 3c–3e, Supplementary Fig. 3a). Identical results have been obtained for reciprocal binding pairs. These data indicate that the first variable Ig domain of sDscamα is sufficient to determine binding specificity.

**Fig. 3.**
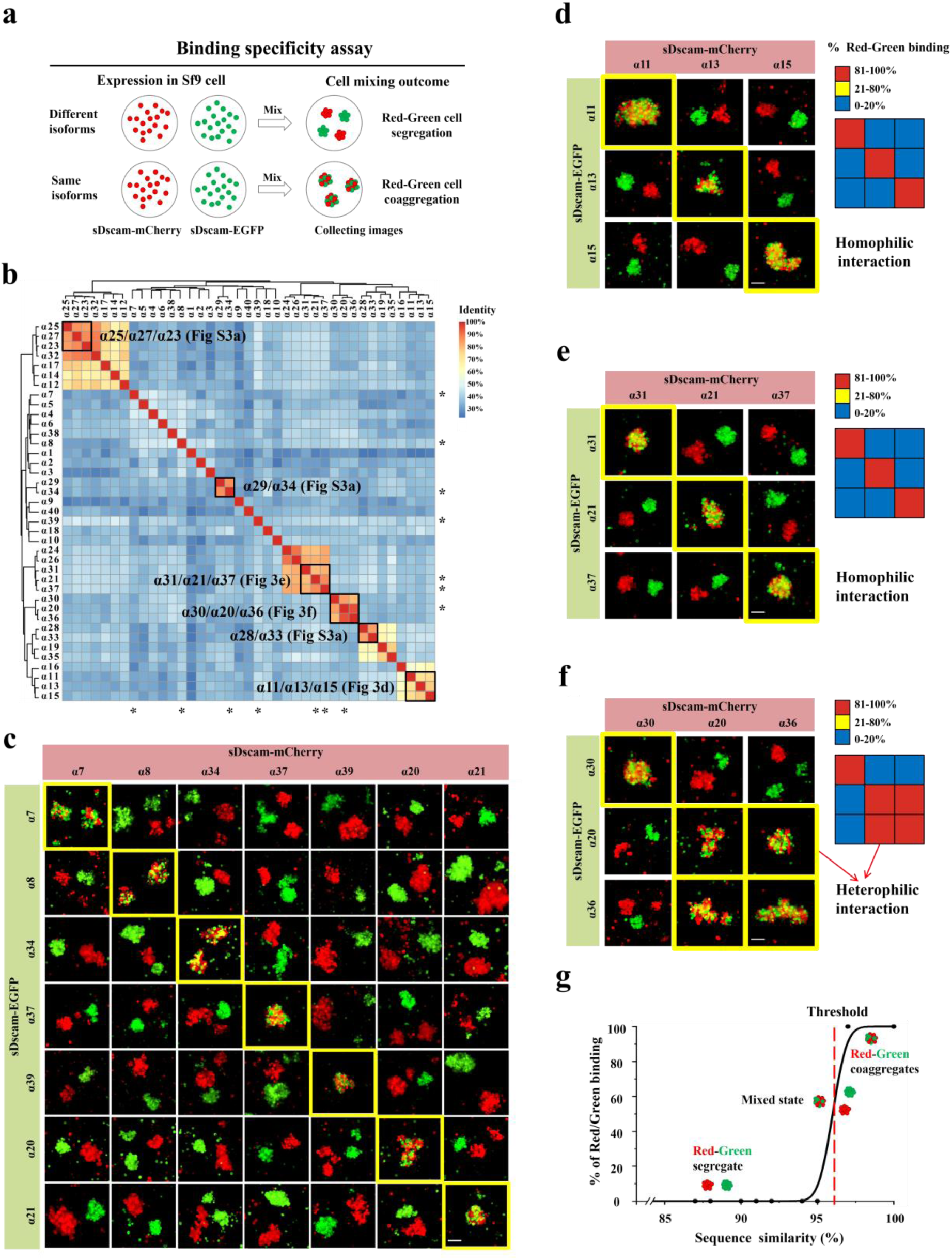
sDscam isoforms engaged in highly specific homophilic interactions. (a) Schematic diagram of the binding specificity assay. Cells expressing mCherry- or EGFP-tagged sDscam isoforms were mixed and assayed for homophilic or heterophilic binding. The outcome of cell aggregation included red-green cell segregation and red-green cell coaggregation. (b) Heat map of pairwise amino acid sequence identities of the Ig1 domains of sDscamα isoforms and their evolutionary relationship. Subsets of the isoforms marked by “*” and within the boxed region were assayed in Fig. 3c–f. See also Supplementary Fig. 3a. (**c-e**) sDscamα isoforms with sequence identity for nonself pairs ranging from 50% to 94% in their Ig domains display strict *trans*-homophilic specificity. Pairwise combinations within representative sDscamαs were assayed for their binding specificity. Scale bar, 100 µm. (f) Pairwise combinations within sDscamα20&30&36 pairs were assayed for their binding specificity. sDscamα36 exhibited strong heterophilic binding to sDscamα20, but did not to sDscamα30. (g) Sequence identity of the Ig1 domains correlated with their binding specificity. Comparative analysis indicates that the thresholds for homophilic and heterophilic binding is set by ∼96% sequence identity within their Ig1 domains between sDscamα pairs. Cell mixing outcome of different sDscamα pairs with sequence identity lower than threshold (∼96%) showed red-green segregation, while red-green aggregation was observed in sDscamα pairwise combinations with sequence identity higher than threshold.

However, heterophilic binding occurred between closely related variable domains. For example, sDscamα20 and sDscamα36 (96.8% sequence identity in their Ig1 domains) showed remarkable heterophilic binding (Fig. 3f). A comparative analysis indicated that the thresholds for homophilic and heterophilic binding were ∼ 96% sequence identity in the Ig1 domains of sDscamα pairs (Fig. 3g). We identified only one example of heterophilic binding among the 35 Ig1 pairs. Thus, the vast majority of the sDscamα isoforms exhibited strict homophilic *trans* binding.

Next, we investigated the specificity of the interactions between sDscamβ isoforms, which differ in their Ig1–2 domains at the N-terminus. We generated pairwise sequence identity heat maps of the Ig1–2 domains and found that only one sDscamβ pair shared more than 90% sequence identity between the Ig1 domains (Supplementary Fig. 3b). Because these isoforms did not support homophilic binding when individually assayed, we were unable to assess their binding specificity. In total, we tested 10 pairs of sDscamβ/β and sDscamα/β, respectively. All of the sDscamβ/β and α/β pairs tested bound strictly homophilically (Supplementary Fig. 3c, d). Taken together with the results of sDscamα analysis (Fig. 3), these observations demonstrated a highly homophilic interaction between sDscamα and sDscamβ isoforms.

### Domain shuffling identifies variable Ig1 as key specificity-determining domains

In contrast to sDscamαs, sDscamβs contain two variable Ig domains at the N-terminus. To identify the variable domain responsible for the specificity of *trans* interactions between different sDscamβ isoforms, we constructed a series of Ig-domain swapping chimeras between sDscamβ5v4 and β5v10 isoforms within the same cluster (Fig. 4a). The two isoforms have 46.7% amino acid sequence identity within their variable Ig1 domains. As a result, a chimeric construct encoding sDscamβ5v4 with its Ig1 domain replaced by that of sDscamβ5v10 no longer interacted with its parent sDscamβ5v4, but interacted with sDscamβ5v10 (Fig. 4a, panels ii and iv, Supplementary Fig. 4b). By contrast, a chimeric construct encoding sDscamβ5v4 with its Ig2 domain replaced by that of sDscamβ5v10 still interacted with its parent sDscamβ5v4, but not with sDscamβ5v10 (Fig. 4a, panels i and iii). Identical results were obtained by Ig domain swapping between sDscamβ5v8 and β5v10 (Fig. 4a, panels v-viii), and sDscamβ5v5 and β5v10 (Supplementary Fig.4c). Therefore, the first Ig domain of sDscamβ is the primary determinant of *trans* interaction specificity.

**Fig. 4.**
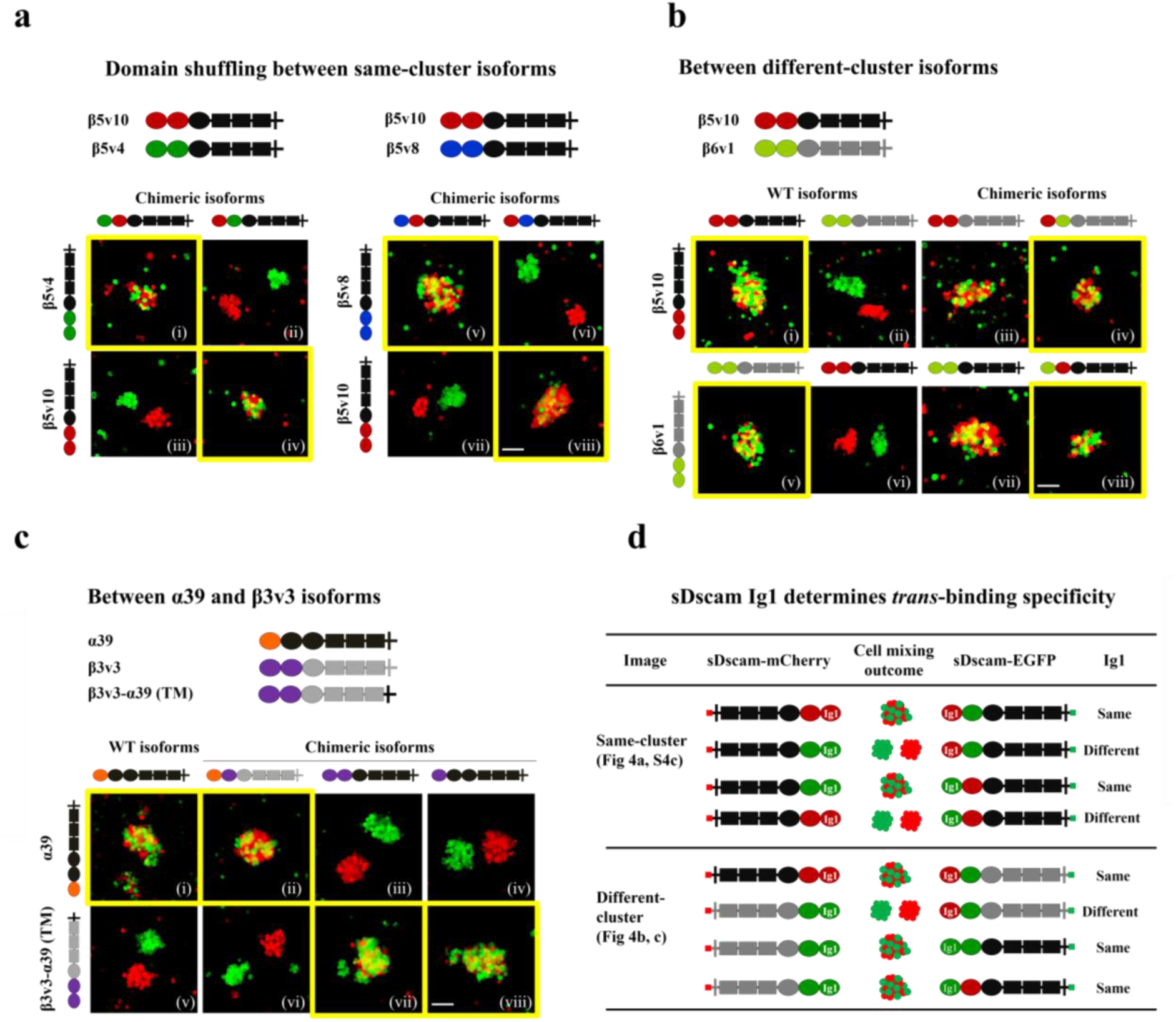
sDscam *trans*-binding specificity is largely dependent on N-terminal Ig1 domain. (a) Domain-shuffled chimeras of sDscamβ5 isoforms and their parental counterparts were assayed for binding specificity. Chimeras in which either the Ig1 or Ig2 domains were replaced with the corresponding domains of cluster-within isoforms swapped or not swapped *trans*-binding specificity. See also Supplementary Fig. 4a, b. (b) Swapped specificity was shown in sDscamβ5v10 and sDscamβ6v1 chimeras. These chimeras were replaced either the Ig1–2 or single Ig1/Ig2 domains with the corresponding domains of different cluster isoforms. See also Supplementary Fig. 4a, b. (c) Domain-shuffled chimeras between sDscamα and sDscamβ3 and their parental counterparts were assayed for binding specificity. See also Supplementary Fig. 4a, b. (d) A summary of schematic of the domain-shuffled sDscam chimeras and their observed binding specificities. These data indicate that the presence of a single common Ig1 domain is essential and sufficient to confer co-aggregation between sDscam isoforms. Scale bar, 100 µm.

Conversely, is a single common Ig1 domain sufficient to confer co-aggregation between sDscamβ isoforms? To determine this, we performed domain swapping between different sDscamβ subfamilies, which differ in both the variable and constant regions (Fig. 4b). A cell aggregation assay indicated that a chimeric construct encoding sDscamβ6v1 with its Ig1 domain replaced by that of sDscamβ5v10 interacted with sDscamβ5v10 (Fig. 4b, panel iv), and *vice versa* (Fig. 4b, panel viii). A similar result has been observed for domain swapping between sDscamα and sDscamβ subfamilies (Fig. 4c). In this case, cells expressing a chimeric construct encoding sDscamβ3v3 with its Ig1 domain replaced by that of sDscamα39 co-aggregated with cells expressing sDscamα39 (Fig. 4c, panel ii), and *vice versa* (Fig. 4c, panel viii). These domain swapping experiments between different sDscam subfamilies indicate that a single same Ig1 domain is sufficient for mediating *trans* binding specificity, at least for the sDscam isoform pairs tested. Conversely, these observations also showed that the constant region of sDscam might not be involved in defining its binding specificity. These data further support a key role for the Ig1 domain in determining the *trans* interaction specificity.

### sDscams interact in *trans* via antiparallel Ig1 self-binding

To gain insight into how the variable Ig1 domain mediates homophilic binding specificity, we carried out homology modeling studies to generate homodimeric complexes of sDscamα Ig1 variants. Based on the crystal structure of variable Ig7 of fly Dscam1 ^37, 38^, we built an Ig1 homodimeric model of sDscamα30 with the SWISS-MODEL program and showed that sDscamα might adopt an antiparallel self-binding fashion of Ig1/Ig1 (Fig. 5a)^39, 40^. Furthermore, docking modeling of each sDscamα revealed that there was a complementary electrostatic potential surface pattern on the ABDE face: positive at one end and negative at the other (Fig. 5a). In this interface docking model, the positive residues might interact with neighboring negative residues to form a salt bridge in the ABDE face. For instance, residue 5 lysine (K) (sDscamα30 Ig1 numbering) in the A strand and residue 12 aspartic acid (D) in the AB loop region are in close structural proximity at the homophilic binding interface, and thus may form a salt bridge (Fig. 5a).

**Fig. 5.**
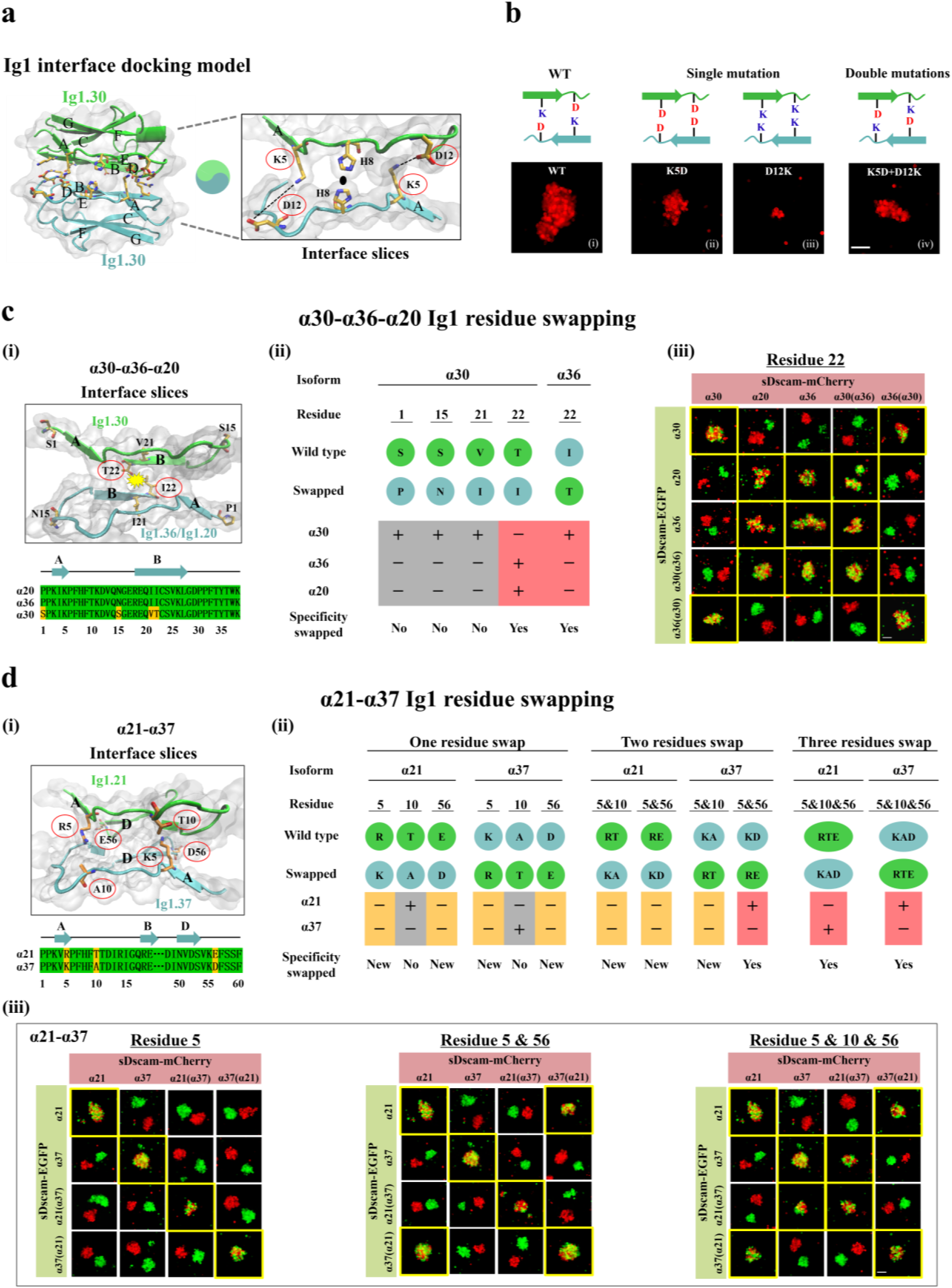
Identification of Ig1 specificity-determining residues. (a) Ig1.30 domain structural modeling. Structural modeling shows that Ig1.30 domain might interact in an antiparallel fashion. Left: The residues represented in licorices (light orange) have been shown a complementary electrostatic potential surface pattern on the ABDE face: positive in one end, and negative in the other end. Right: Slices of the Ig1.30-Ig1.30 interface between strand A subunits. Potential neighboring interact residues (K5 and D12) were shown in licorices (light orange). (b) The single disrupting and double complementary mutation of these candidate residues were assayed for cell aggregation. The double mutation partially rescued the reduced cell aggregates by single point mutations, supporting the antiparallel binding fashion. (c) Residue swapping between sDscamα20&α30&α36 to assess specificity-determining residues. Left: Ig1 docking model and sequence alignments of shuffled regions are shown on the panel i. Four candidate specificity-determining residues (light orange) were located on adjacent B strands. Middle: Panel ii shows schematic representation of residue swapping mutants used in the experiments, along with a summary of results from binding specificity. Right: Panel iii shows the binding specificity of isoforms containing wild-type and swapped residue 22. Swapping of residue S1, S15, and V21 in Ig1.30 to Ig1.36 did not swap *trans*-binding specificity (data not shown), while residue 22 swapping between Ig1.30 and Ig1.36 swapped binding specificity. See also Supplementary Fig. 5a. (d) Residue swapping of variable Ig1 between sDscamα21 and α37. Left panel (i) shows Ig1 docking model and sequence alignments of shuffled regions. Three candidate specificity-determining residues (light orange) were located on adjacent A and D strands. Right panel (ii) shows the schematic diagrams of residue swapping mutants used in the experiments, along with observed binding specificity. Lower panel (iii) shows cell aggregation assays of isoforms containing wild-type and residue-swapped Ig1 domains. Swapping of either one of three residues between in Ig1.21 to Ig1.37 did not swap binding specificity, and swapping of two of three residues between in Ig1.21 to Ig1.37 partially swapped binding specificity, and swapping of all three residues between in Ig1.21 to Ig1.37 fully swapped binding specificity. See also Supplementary Fig. 5g. Scale bar, 100 µm.

To confirm this, we performed single and double complementary mutations of these candidate residues and assessed their ability to mediate cell aggregation. As a result, a single K mutation of residue 5 in the A strand or a D mutation of residue 12 in the AB loop region weakened cell aggregation (Fig. 5b, panel ii and iii). These results indicate that residue K5 in the A strand and residue D12 in the AB loop region are necessary for efficient self-recognition of sDscamα30. Interestingly, double K/D mutants (K5D; D12K), which are thought to reform a salt bridge at the interface, partially restored cell aggregation (Fig. 5b, panel iv), validating an antiparallel self-binding mode of Ig1/Ig1. Therefore, the Ig1 domains of sDscamα adopt an antiparallel self-binding conformation.

### Identification of sDscam Ig1 specificity-determining residues

We next identified the Ig1 specificity-determining residues by structural modeling and mutagenesis studies. Candidate specificity-determining residues were predicted using the closely related Ig1 of sDscam (Supplementary Fig. 5a). Because the homophilic binding specificity of fly Dscam1 Ig7, a Chelicerata sDscam Ig1 homologue, is largely determined by electrostatic and symmetry axis residues ^37, 38^, different residues between each pair were screened based on their electrostatic and shape complementarity. A remarkable example is the closely related α20/α36/α30 pairs, in which sDscamα36 exhibited robust heterophilic binding to sDscamα20 but no heterophilic binding to sDscamα30 (Fig. 3f). Four amino acids were different between the Ig1 domains of α30 and α36; however, only residue 22 of the B sheet, which resided at the symmetry axis, was predicted to influence shape complementarity (Fig. 5c, panel i). Therefore, we speculate that this residue is a candidate specificity determinant.

To confirm this, we swapped them between isoforms and examined the binding specificities of the swapped isoforms and their parents (Fig. 5c, panel ii). Swapping residue 22(T/I), but not the other three residues (1S/P, 15S/N, and 21V/I), switched the *trans* binding specificity between sDscamα30 and sDscamα36 (Fig. 5c, panel ii and iii). Cells expressing a swapped sDscamα30 isoform in which residue 22T was replaced with 22I from sDscamα36 and sDscamα20 intermixed with cells expressing sDscamα36 and sDscamα20 (Fig. 5c, panel iii). By contrast, these cells segregated from those expressing the parent isoform from which the swapped residue derived, and *vice versa*. Thus, a single residue is sufficient to determine binding specificity in some sDscamα pairs.

However, swapping residue 22(I/T) of Ig1 between sDscamα11 and α15, or between sDscamα13 and α15, did not swap binding specificity but instead produced novel homophilic binding specificity (Supplementary Fig.5c). These observations suggest that additional residues at the Ig1/Ig1 interface also contribute to binding specificity. By swapping interface residues that differed between other pairs of Ig1 variants (sDscamα11 and α13; α23 and α27), the symmetry axis residue 52 (M/V; L/V) of the D sheet was shown to alter, but not swap, the binding specificity (Supplementary Fig.5e). In the latter case (α27/α23), separate green and red homophilic aggregates are formed, but they now adhere to one another (Supplementary Fig.5e, panel 18 and 20). By contrast, swapping residue 26 (N/I) between sDscamα11 and α13, which was neither electrostatic nor located at the symmetry axis, did not alter the binding specificity (Supplementary Fig.5d). Collectively, these data indicate that residues 22 of the B sheet and 52 of the D sheet act as specificity determinants, at least for some pairs of sDscam isoforms.

Another representative example of specificity swapping is shown for the α21/α37 pair (Fig. 5d). In the α21/α37 pair, three different residues (5, 10&56) were filtered as candidate specificity determinants, one of which interacted at the antiparallel self-binding interface by double complementary mutations (Fig. 5b). We found that swapping one of the three different residues (5, 10&56) between sDscamα21 and α37 did not swap binding specificity (Fig. 5d, panel iii, Supplementary Fig. 5g). By contrast, swapping of all three residues of sDscamα21 to those of sDscamα37 swapped the binding specificity (Fig. 5d, panel iii). Conversely, swapping two of the three residues partially or completely altered the binding specificity between sDscamα21 and α37 (Fig. 5d, Supplementary Fig. 5g). Similarly, swapping of all three residues 6, 19 and 52 between sDscamα23 and α27 fully switched the binding specificity (Supplementary Fig. 5f). Therefore, these residues are involved in determining binding specificity. This may also, at least in part, mirror Ig1 diversification during sDscam evolution, where initial mutations after exon duplication could lead to promiscuous binding and additional combinatorial mutations could eventually produce a highly specific homophilic interaction distinct from their parents.

In summary, by structural modeling and mutagenesis studies, we identified specificity-determining residues for sDscam isoforms (Supplementary Fig. 5b). These residues frequently resided at the center or side-chain of symmetric homo-dimerized interfaces, which exhibit electrostatic and shape complementarity between the ABED interface strands. As the six pairs of variable Ig domains used in these residue swapping experiments are not closely related (Supplementary Fig.5a), these interface residues likely contribute to determining homophilic specificity for other sDscam isoforms as well.

### sDscams form high order *cis*-multimers independent of their *trans* interactions

Having established that sDscam isoforms are mediated by an Ig1/Ig1 antiparallel self-binding between apposed cell surfaces, we attempted to explain how sDscam isoform diversity mediates self-recognition in Chelicerata. As estimated by the number of Ig1 or its orthologues, the number of Dscam isoforms is in the range of ∼ 100 across the Chelicerata species ^32^, at least two orders of magnitude lower than that in flies. Moreover, as shown above (Fig. 1), almost half of sDscam isoforms, when expressed individually, failed to engage in homophilic interactions in the cell aggregation assay (Fig. 1d). Therefore, the small number of sDscam isoforms in Chelicerata alone cannot account for its non-self-discrimination as for the homophilic interaction model in *Drosophila* Dscams. Thus, how do these nonclassical sDscam isoforms mediate homophilic interactions? Given the striking organizational resemblance between the scorpion *sDscam*s and mammalian *Pcdh*s, we speculate that via *cis*-multimers scorpion sDscams function as vertebrate-clustered Pcdh isoforms ^22, 23, 29^.

To confirm this hypothesis, we first performed coimmunoprecipitation (co-IP) analyses to investigate the interactions between sDscam isoforms *in vitro* (Fig. 6a). When HA-sDscamβ6v2 and Myc-sDscamβ6v2 are coexpressed by coinfecting with individual recombinant viruses, Myc-tagged proteins could strongly coimmunoprecipitate with HA-β6v2 (Fig. 6b). Further co-IP experiments indicate that sDscam proteins from different subfamilies tested interacted strongly with each other (Fig. 6b, Supplementary Fig. 6a), exhibiting no specificity between different isoforms. Because sDscamαΔIg1ΔTM could interact with other sDscamΔIg1 or sDscamΔIg1-2 mutants (Supplementary Fig. 6b), in which Ig1-domain deletion has ablated homophilic *trans* interactions, robust co-IP between HA-sDscam and Myc-sDscam should not result from *trans* interactions, but *cis* interactions. These results demonstrate that sDscams interacted with each other in *cis* with almost no specificity between different isoforms.

**Fig. 6.**
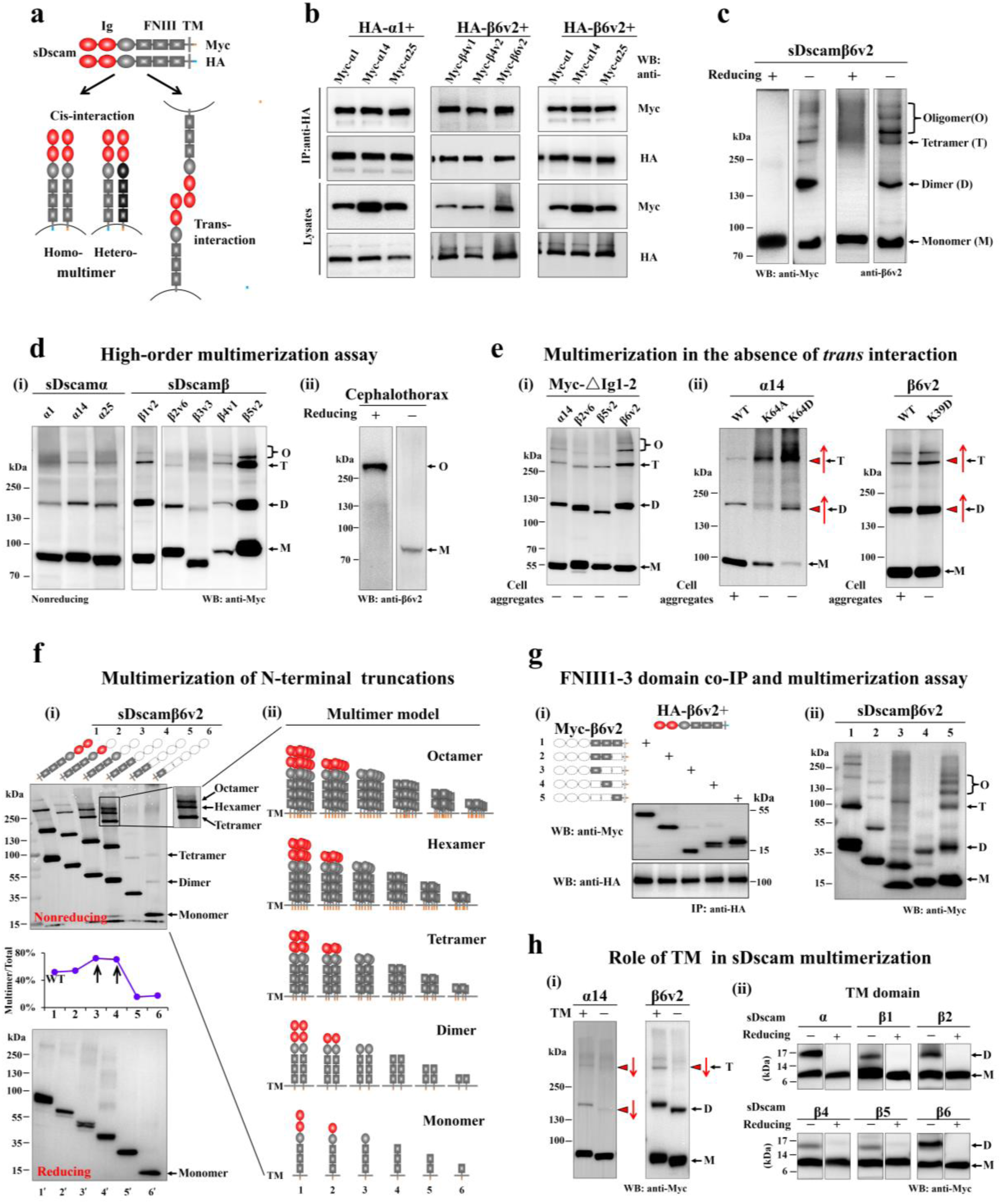
sDscams form high order *cis*-multimers mediated by FNIII1–3 and TM domains. (a) Schematic of *cis*- and *trans*-interaction of sDscam. sDscam monomers interacted in a parallel fashion to form a homomultimer or heteromultimer complex, while *trans*-multimers are formed between two opposing cells in an antiparallel fashion. (b) All sDscamα and sDscamβ isoforms tested interacted strongly with each other in co-IP experiments. Lysates from Sf9 cells cotransfected with sDscamα1, sDscamβ4v1 and sDscamβ6v2 bearing a C-terminal HA-tag (HA-α1, HA-β4v1 and HA-β6v2) and different Myc-tagged sDscam isoforms were immunoprecipitated using anti-HA antibody and probed with anti-Myc or anti-HA antibodies. See also Supplementary Fig.6a. (c) sDscamβ6v2 expressed in Sf9 cells formed *cis*-multimers. Lysates from Sf9 cells expressing sDscamβ6v2 were run on SDS-PAGE in the presence of nonreducing or reducing agents, and analyzed by western blot with Myc antibody (lane 1-2), and with sDscamβ6v2 antibody (lane 3-4). (d) Multimerization assay of sDscam expressed in Sf9 cells (panel i) and from the scorpion cephalothorax (panel ii). Lysates from the scorpion cephalothorax were resolved on a SDS/PAGE gel under the nonreducing or reducing conditions, and analyzed by western blot with sDscamβ6v2 antibody (panel ii). (e) sDscams formed high order *cis*-multimers in the absence of *trans* interaction. (i) Proteins lacking Ig1-2, which have ablated homophilic *trans-*interactions, was able to form robust multimers. (ii) Single residue mutations (e.g., α14K64A, α14K64D and β6v2K39D), which ablated homophilic cell aggregation, caused increased multimerization. (f) A series of N-terminal truncations of the extracellular domain of sDscamβ6v2 fused with Myc-tag were examined for multimerization assay. Nonreducing (upper panel i) or reducing (lower panel i) SDS/PAGE gels were analyzed, with graph of the ratio change of multimer/total below nonreducing gel. Schematic architecture of *cis*-multimer was depicted on the right (panel ii). See also Supplementary Fig.6e. (g) Co-IP and multimerization assay of FNIII1–3s. sDscamβ6v2 interacted strongly with each truncated protein expressing individual or combined domain of FNIII1–3s (panel i), and each truncated protein could form strong *cis-*multimers (panel ii). (h) Transmembrane domain promotes the formation of sDscam *cis*-multimers. TM domain deletion strikingly reduced sDscam multimerization (Downward red arrow, panel i). The TM peptides expressed from various sDscams could dimerize strongly (panel ii). See also Supplementary Fig.6g.

To further characterize the *cis* interaction between sDscam isoforms, we performed multimer analysis by western blotting in the absence or presence of reducing agents. In reducing SDS/PAGE gels, sDscamβ6v2 migrated with a single molecular weight of ∼ 80 kDa, which corresponded to the size of the monomer (Fig. 6c). Under non-reducing conditions, however, several large bands migrated behind the monomer, which corresponded to the size of sDscam assembly of putative dimer, tetramer, and larger multimers (Fig. 6c). Notably, all sDscam assembly sizes were a multiple of the dimer, suggesting that the dimer acts as a basic recognition unit for sDscam, and clusters into tetramer and higher order oligomeric complexes.

Similarly, we performed multimerization analysis by expressing proteins of all other sDscamα and sDscamβ1–β5 subfamilies. We observed multimerization to different extents in all sDscam proteins investigated, suggesting that sDscam proteins from different subfamilies are able to homo-multimerize (Fig. 6d, panel i). In these cases, we found that individual sDscam isoforms from the same cluster (i.e., sDscamα1, α14, α25) exhibited largely similar multimerization behavior, which varied markedly among the sDscam proteins from different subfamilies. Furthermore, we have observed abundent endogenous sDscamβ6v2 multimers in the cephalothorax of scorpion (Fig. 6d, panel ii). Taken together, our results demonstrate that sDscams can form strong multimers *in vivo*.

It is possible that sDscam multimers result from *trans* interactions in an antiparallel orientation (Fig. 6a). However, sDscamβ1v2 and β4v1, which did not show homophilic *trans* interactions in cell aggregation assays, still exhibited robust multimers (Fig. 6d, panel i). Moreover, Ig1–2 domain deletions, which ablated homophilic *trans* interactions, still exhibited strong multimerization (Fig. 6e, panel i). Therefore, sDscam *cis* interactions occurred even in the absence of *trans* interactions. To preclude the effect of sDscam size on multimerization, we performed single amino acid mutations in the Ig1 domain of sDscams and examined the effect on multimerization (Fig. 6e, panel ii). Single mutations, which disrupted homophilic *trans* interactions in a cell aggregation assay, heightened multimerization (Fig. 6e, panel ii). These results support the notion that sDscams multimerized in *cis* independently of their *trans* interactions and indicate that robust sDscam multimers result from *cis* interactions.

### sDscam *cis* multimerization is mediated by the extracellular FNIII1**–**3 domains

To elucidate how sDscams form high order *cis*-multimers, we constructed a series of truncations of the extracellular domain of sDscamβ6v2 fused with a Myc-tag for *cis* multimerization assay by sequentially deleting the domain from the N-terminus (Fig. 6f). We found that the truncated proteins lacking Ig1 (β6v2ΔIg1), Ig1–2 (β6v2ΔIg1–2), and Ig1–3 (β6v2ΔIg1–3) exhibited robust multimerization (Fig. 6f, panel i, lane 2–4). Notably, the truncated proteins lacking Ig1, Ig1–2, and Ig1–3 exhibited multiple bands, which corresponded to the size of sDscam assembly putative dimers, tetramer, hexamer, octamer, and larger multimers as the entire extracellular domain, respectively (Fig. 6f, panel i). Therefore, the absence of Ig1, Ig1–2, and Ig1–3 did not influence the assembly pattern of multimerization. Importantly, based on the unaltered assembly pattern of multimers in a series of successively truncated proteins, we reason that sDscams are assembled into *cis*-dimer and higher-order multimeric complexes in a parallel orientation (Fig. 6f, panel ii). Furthermore, sDscam mutants lacking Ig1–2 or Ig1–3 exhibited the most heightened *cis* multimerization (Fig. 6f, arrows in panel i). Together, these data strongly suggest that the Ig1–3 domains of sDscam are dispensable for *cis* interactions, and that membrane-proximal FNIII1–3 domains are sufficient for efficient *cis* multimerization.

However, when reduced to one or two membrane-proximal FNIII domains, the resulting constructs (i.e, β6v2ΔIg1–FNIII1 and β6v2ΔIg1–FNIII2) exhibited markedly reduced multimerization (Fig. 6f, lane 5, 6). Because these constructs contained the same transmembrane domain, the discrepancy between them is likely due to the presence of different FNIII domains. To further identify the extracellular domains of sDscam that contribute to *cis* multimerization, we constructed mutants encoding a single or two continuous extracellular domains with an HA tag and co-transfected them with sDscamβ6v2 (Fig. 6g). Co-IP experiments revealed that the deletion constructs containing individual or multiple continuous FNIII domains were capable of binding to sDscamβ6v2 (Fig. 6g, panel i, Supplementary Fig. 6c), indicating that sDscamβ6v2 can interact with each FNIII domain. This observation was supported by *cis* multimerization assays, which showed that each of the truncated β6v2 proteins could form strong *cis*-multimers (Fig. 6g, panel ii). These data indicate that *cis* multimerization is jointly mediated by all three membrane-proximal FNIII1–3 domains. This result is also consistent with computational modeling using the ZDOCK server ^41^, by which sDscamβ6v2 could form a homodimer via multiple parallel interfacial regions involving the FNIII1–3 domains (Supplementary Fig. 6d). Because the sDscamβ6v2 FNIII1–3 domains lack cysteine residues, they likely mediate *cis* multimerization by noncovalent mechanisms.

We subsequently carried out multimerization assays of all other sDscamα and sDscamβ1– β5 proteins by truncating the extracellular domain as for sDscamβ6v2. Representative examples of multimerization truncation are shown for sDscamα14, sDscamβ1v2, sDscamβ2v6, and sDscamβ5v2 in Supplementary Fig. 6e. Consistent with sDscamβ6v2, the absence of the N-terminal Ig1–3 domains increased the extent of *cis* multimerization in almost each sDscam (arrows in graph of Supplementary Fig. 6e). These observations suggest that the membrane-proximal FNIII1–3 domains of sDscams are sufficient for efficient *cis* multimerization. However, when deleting the FNIII domains, individual sDscams from different subfamilies exhibited different patterns of alteration of multimerization (Supplementary Fig. 6e). For instance, the FNIII2–3 peptide (i.e, β5v2ΔIg1–FNIII1) exhibited similar multimerization to the FNIII1–3 peptide (β5v2ΔIg1–3) in sDscamβ5v2 (Supplementary Fig. 6e, panel iv, lane 4, 5), while the FNIII2–3 peptide (i.e, β6v2ΔIg1–FNIII1) exhibited much less multimerization than the FNIII1–3 peptide (β6v2ΔIg1–3) in sDscamβ6v2 (Fig. 6f, panel i, lane 4, 5). This suggests that the contribution of individual FNIII domains to *cis* interactions differs markedly among sDscams. Taken together, our results indicate that all three membrane-proximal FNIII1–3 domains engage in *cis* multimerization of sDscams.

In summary, our data reveal that all three membrane-proximal FNIII1–3 domains engaged in high order *cis* multimerization, while the first N-terminal Ig1–3 domains were dispensable for *cis* interactions in all sDscams investigated. Notably, despite their general promiscuity, all FNIII1–3 domains of sDscam may contribute to *cis* interactions, with the relative contributions of each domain to *cis* multimerization varying from one sDscam cluster to another.

### Transmembrane domain promotes the formation of sDscam *cis*-multimers

We subsequently investigated whether and how the transmembrane (TM) domain of sDscam contributes to high-ordered *cis* multimerization. To this end, we first examined how TM deletion affected self-multimerization of sDscam. Each of the truncated proteins lacking the TM domain was capable of multimerizing, indicating that the extracellular domain is sufficient to confer efficient *cis* multimerization (Fig. 6h, Supplementary Fig. 6f). However, the multimerization efficiency was markedly reduced in most sDscamΔTM mutants (arrows in Fig. 6h, Supplementary Fig. 6f). For example, dimer and tetramer were reduced in sDscamαΔTM, while tetramer was almost undetectable in β6v2ΔTM (Fig. 6h, panel i). Overall, these data demonstrate that although the extracellular region alone might engage in the formation of multimers *in cis*, the presence of the transmembrane domain greatly enhances the accuracy and efficiency of sDscam *cis* multimerization.

To further determine the role of the TM domain in *cis* multimerization, we examined whether sDscam transmembrane peptides, in the absence of regulation by soluble domains, have the propensity for self-multimerization. When the expressed transmembrane peptides were treated in the absence or presence of reducing agents, we observed dimerization of the transmembrane peptides expressed from all six sDscams investigated (Fig. 6h, panel ii), while the addition of reducing agents precluded dimer formation. These observations indicate that the presence of the transmembrane domain alone is sufficient to confer efficient *cis* dimerization in each sDscam. However, we have not observed higher order multimers (i.e., tetramer, hexamer) for sDscam transmembrane peptides as extracellular proteins. Notably, although individual transmembrane peptides from all sDscams engaged in dimerization, the efficiency of dimerization varied remarkably among them, with the greatest efficiency for sDscamα and sDscamβ6 and the least for sDscamβ4 and sDscamβ5 (Fig. 6h, panel ii). By contrast, TM peptides from sDscamβ4 and sDscamβ5 accounted for less than 10% of dimers (Fig. 6h, panel ii). These results further implicate the transmembrane domain in sDscam *cis* multimerization.

To further explore how the transmembrane domain promotes sDscam *cis* multimerization, we analyzed the roles of transmembrane residues using SWISS-MODEL ^39, 40^. As expected from sequence-based α-helical transmembrane prediction tools, several of the residues in the membrane form a contiguous helix in sDscamβ6v2, which have a propensity to dimerize via covalent or noncovalent interactions (Supplementary Fig. 6g, panel i). Notably, the transmembrane of some sDscams (i.e, sDscamβ2, β3), which did not contain cysteine residues, exhibited considerable dimerization (Fig. 6h, panel ii). This observation suggests that disulfide bonds within the transmembrane domain are not essential for *cis* dimerization at least in some sDscams. A further mutation experiment showed that cysteine residue mutation in the transmembrane domain of sDscamβ6v2 did not markedly affect the formation of *cis*-multimers in both the β6v2 and β6v2ΔIg1–3 constructs (Supplementary Fig. 6g, panel ii). Collectively, these results demonstrate that the transmembrane domain of each sDscam mediated *cis* multimerization, likely via a noncovalent mechanism.

### Coexpression of multiple sDscam isoforms diversify homophilic specificities

Finally, because all sDscamα isoforms and some of sDscamβs mediated homophilic *trans* bindings (Fig. 1d) and interacted with each other in *cis* without specificity (Fig. 6), we tested how combinatorial expression of multiple sDscam isoforms diversified binding specificities. In all cases, cells coexpressing two sDscamα combinations failed to coaggregate with cells expressing two different sDscamα combinations. However, intermixed cell aggregates of cells that coexpressed identical sDscamα combinations were observed (Fig. 7a). Consistent with these observations, co-IP experiments showed different sDscamαs interacted strongly with each other when coexpressed (Supplementary Fig. 7b, lane1-2). Similar data were obtained for each of the sDscamα/β pairs (Fig. 7b, Supplementary Fig. 7b, lane 3-4). These results suggest that homophilic specificity by coexpressing two distinct isoforms depends on the identity of the isoform pair but not a single isoform. These results strongly suggest that sDscams interact *in cis* so as to create new homophilic specificities that differ from the specificities of the individual sDscam isoforms.

**Fig. 7.**
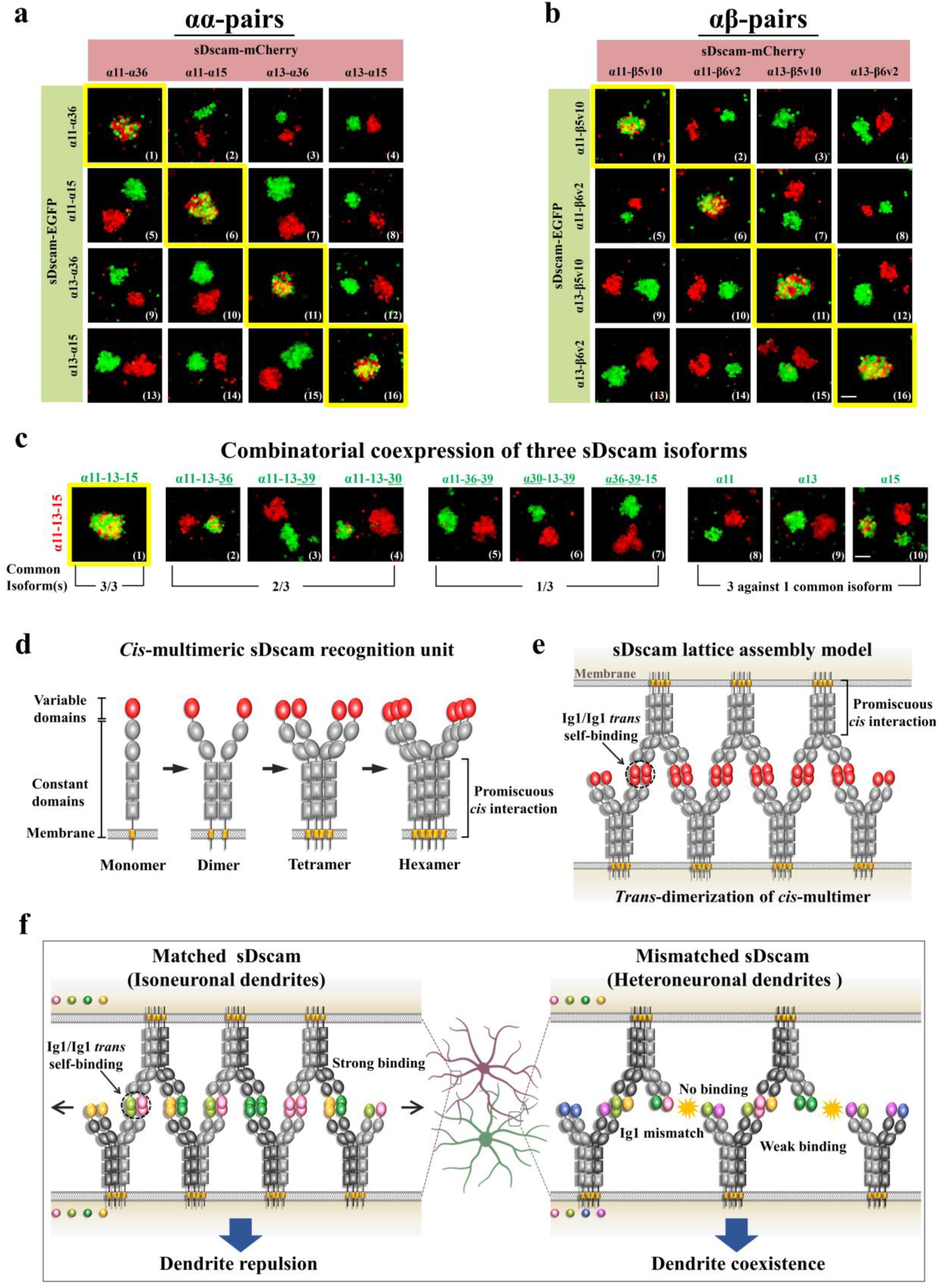
The model for sDscam-mediated cell-cell recognition. (**a, b**) Combinatorial coexpression of multiple sDscam isoforms generates unique cell surface identities. Cells coexpressing an identical or a distinct set of sDscamα (**a**) and sDscamβ5–β6 isoforms (**b**) were assayed for coaggregation. See also Supplementary Fig.7a. (c) Cells coexpressing three distinct mCherry-tagged sDscam isoforms were assayed for interaction with cells expressing an identical or a distinct set of GFP-tagged sDscam isoforms. The nonmatching isoforms between two cell populations are underlined. See also Supplementary Fig.7a. Scale bar used in (a–c), 100 μm. (d) Model of *cis*-multimeric sDscam recognition units. The *cis* interface is located on long-range region from FNIII1 to TM domains. Since all sDscam assembly sizes were the fold of dimer, we suggest that dimer might act as basic recognition units for sDscam, and then clustered into tetramer and higher orderly oligomeric complexes. (e) The model for sDscam-mediated cell-cell recognition. In this model, each sDscam *cis*-tetramers could interact multiple *cis*-tetramers on apposed cell surfaces via independent *trans* Ig1/Ig1 self binding, thereby forming a connected latticed assembly of proteins between cells. (f) The model of sDscam-mediated neuronal self-recognition and self–non-self discrimination. Due to identical sDscam isoforms in two neurites of the same neuron, the *trans*-dimerization of *cis*-multimers could lead to a dense and connected lattice assembly between two apposing cell surfaces, thus triggering strong homophilic interactions and then inducing neurite repulsion. In contrast, mismatched Ig1s between two neighbouring neurons lead to a scattered or sparse connected lattice assembly between apposing cell surfaces, triggering weak homophilic interactions. Thus, the resulting downstream signaling is below the threshold level and fails to initiate neurite repulsion.

To further examine combinatorial homophilic specificities, we coexpressed distinct sets of three sDscam isoforms, and then evaluated their ability to coaggregate with cells containing various numbers of mismatches (Fig. 7c). In all cases, only cells expressing identical isoform combinations formed intermixed coaggregates, while cells expressing mismatched isoforms displayed separate red and green aggregates (Fig. 7c). Remarkably, even a single isoform common between cells coexpressing three isoforms and expressing one isoform caused separate red and green aggregates (Fig. 7c, panels 8-10). Taken together, these results strongly suggest that sDscam isoforms evolved a unique combinatorial multimerization to diversify homophilic specificities.

## Discussion

Here, we provide compelling evidence that different combinations of sDscam isoforms interact *in cis* to significantly expand homophilic *trans* recognition specificities in Chelicerata. Specifically, we showed that sDscam isoforms form promiscuous *cis*-multimers involving the membrane-proximal FNIII1–3 and transmembrane domains at the cell surface that associate specifically in *trans* via Ig1/Ig1 self-binding. Thus, our data revealed that Chelicerata sDscams mediated highly isoform-specific homophilic interactions via a “hybrid” mechanism between fly Dscam1 and vertebrate Pcdhs. Below, we discuss the molecular basis of the homophilic interactions of sDscam isoforms, and the implications of sDscam architecture in neuronal self-recognition and non-self-discrimination, with particular emphasis on comparison to fly Dscam1 and vertebrate Pcdhs.

### Chelicerata sDscams form high order *cis*-multimers

Our data clearly indicate that sDscam isoforms could form robust *cis*-multimers even in the absence of *trans* interactions (Fig. 6). This is supported by several lines of independent evidence that distinct sDscam isoforms from different clusters can be coimmunoprecipitated (Fig. 6b, Supplementary Fig. 6a), and that sDscams are present in high-molecular-weight, detergent-solubilized assembly complexes from scorpion cephalothorax (Fig. 6d, panel ii). In addition, we observed altered recognition specificity when multiple sDscam isoforms were coexpressed (Fig. 7a-c), further supporting their *cis* multimerization. Such *cis* multimerization of Chelicerata sDscams is in sharp contrast to that of fly Dscam1, which appear interacts as *cis* monomers. Although the individual orthologue of FNIII1–3 domains in Chelicerata sDscams is present at the membrane-proximal region of fly Dscam1, the overall structure of the membrane-proximal region might be reconstituted in evolutionary transitions from classical Dscam to shortened sDscams. Because sDscam originated from classical Dscam after Chelicerata speciation ^14, 33^, sDscam *cis* multimerization might be in large part the result of evolutionary co-adaptation of these membrane-proximal domains as a *cis* interaction interface for combinatorial recognition specificity.

Although Chelicerata sDscams form *cis*-multimers like Pcdh isoforms, they differ in at least two major aspects. Firstly, Chelicerata sDscams form *cis*-multimers via a *cis* interaction interface distinct from Pcdhs. The latter form *cis* dimers mediated by membrane-proximal EC6 or both EC5 and EC6 ^23,24,25, 29^. By contrast, the formation of sDscam *cis*-multimers might be mediated by combining membrane-proximal FNIII1–3 and transmembrane domains (Fig. 6f-h). Although individual recognition sites located in the extracellular and transmembrane domains alone might engage in the formation of *cis*-multimers, a long-range *cis* interface encompassing the FNIII1 to transmembrane domains could assure the accuracy and efficiency of *cis* multimerization. Thus, the sDscam *cis* interface involves a much larger proportion of the extracellular region than vertebrate Pcdhs. This discrepancy in the *cis* interface may at least in part account for the markedly different *cis* multimerization ability of Pcdhs and sDscams, whereas a γ-Pcdh lacking EC1–3 domains interacted weakly ^22^ while sDscams lacking Ig1–3 domains exhibited robust *cis* multimerization for the one sDscamα and six sDscamβ subfamilies tested (Fig. 6, Supplementary Fig. 6).

Furthermore, sDscams formed more complex *cis*-multimers than Pcdhs. Although γ-Pcdhs reportedly form *cis* tetramers ^22^, it is generally recognized that Pcdhs form *cis*-dimers, as evidenced by dimerization of truncated Pcdhs lacking the EC1 or EC1–2 domain in solution ^25, 29^. By contrast, our evidence showed that sDscams assembled into *cis*-dimer, tetramer, and even higher-order oligomeric complexes (Fig. 6, Supplementary Fig. 6). Therefore, sDscams seem to exhibit more efficient and complex *cis* multimerization than Pcdhs, presumably due to their larger interaction interface.

### sDscams mediate highly specific homophilic recognition via self-binding variable Ig1

We showed that all sDscamα isoforms can mediate homophilic interactions (Fig. 1d, 3b-f, Supplementary Fig. 3a), while a majority of sDscamβ1–β6 isoforms does not interact homophilically (Fig. 1d). However, sDscamβ chimeric constructs (i.e, sDscamβ4v1) produced by deleting or replacing their partial constant region mediated homophilic interactions (Fig. 2c, panels xv and xvi; Supplementary Fig. 2a, panel 15-17; Fig. 2f, panel i and ii), indicating that the failure in homophilic recognition is not due to incompatibility of the *trans* self-binding interface. Thus, it seems likely that at least some of these sDscamβs mediate self-recognition, although the mechanism is not yet understood. Alternatively, because sDscam *cis*-multimer assembly resembles the structures of antibody and T-cell receptor (TCR) in vertebrate, both comprising N-terminal variable Ig domains and a C-terminal constant domain, it is likely that scorpion sDscam isoforms participate in heterophilic binding with other proteins or pathogens as vertebrate antibody and TCR. Moreover, ∼ 100 sDscam isoforms in scorpion could potentially form a repertoire of 10^8–12^ structurally variable assemblies, which is compatible with the order of magnitude of the diversity of antibodies in vertebrates. Notably, pancrustacean Dscam1 isoforms play an immune-protection role in bacterial challenge ^42,43,44^. Given the extraordinary diversity and high structural similarity between Chelicerata sDscam and vertebrate antibody and TCR, it is attractive to speculate that sDscam diversity plays a role in invertebrate immunity.

Based on structural modeling and mutagenesis experiments, we demonstrate that sDscam homophilic specificity is determined by an antiparallel Ig1/Ig1 self-binding (Fig. 5a, b). Because the Ig1 domain of Chelicerata sDscams is orthologous to the Ig7 of fly Dscam1, it is reasonable that they have an identical antiparallel self-binding architecture. Indeed, site-directed swapping mutagenesis revealed that Ig1 of Chelicerata sDscams shared several key specificity-determining residues with Ig7 of fly Dscam1 isoforms^37, 38^ (Fig. 5c, d, Supplementary Fig. 5). However, the homophilic specificity of fly Dscam1 is determined via three independent antiparallel self-binding modules, Ig2/Ig2, Ig3/Ig3, and Ig7/Ig7 (Fig. 8) ^37, 38^, which can assemble in different combinations to generate a repertoire of tens of thousands of self-binding interfaces. By contrast, Ig1/Ig1 self-binding of Chelicerata sDscams only generated ∼ 100 distinct *trans* interfaces. If Chelicerata sDscams use the same neuronal self-recognition mechanism as fly Dscam1, this number is not sufficient to discriminate self from non-self ^11^. Therefore, *trans* homophilic interactions in Chelicerata sDscams likely proceed via a mechanism distinct from that of fly Dscam1.

**Fig. 8.**
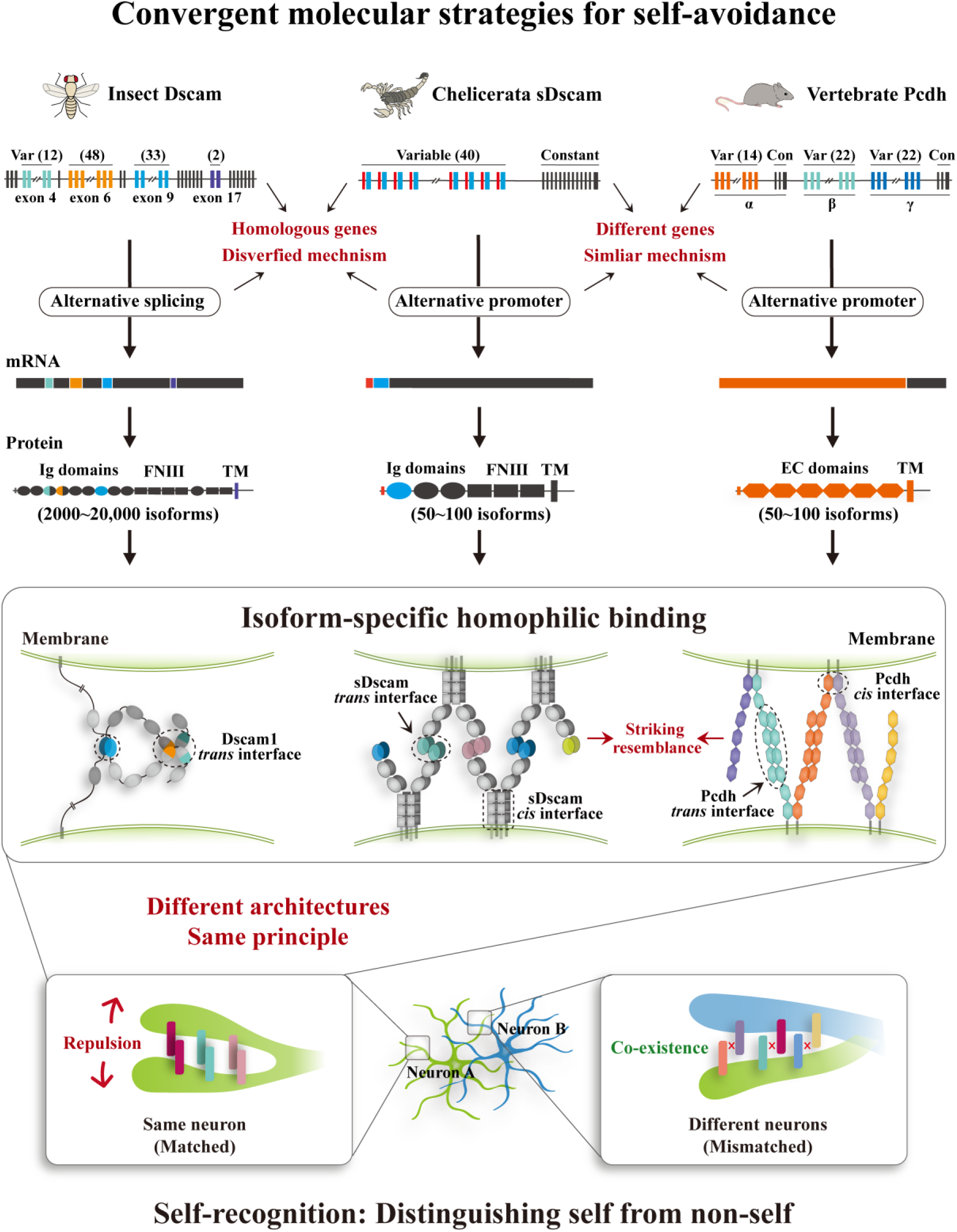
Chelicerata sDscams show more parallels with vertebrate Pcdhs than *Drosophila* Dscam1. Chelicerata sDscams show more parallels with vertebrate Pcdhs than *Drosophila* Dscam1. *Drosophila Dscam1* generates ten thousands of protein isoforms through alterntive splicing^9^. Chelicerata *sDscam* genes employ alternative promoter to generate extensive isoforms as vertebrate Pcdhs ^17, 33^. Three of them encode a large number of single-transmembrane protein isoforms, and the encoded proteins engage in isoform-specific homophilic binding. This suggests that different phyla seem to have used different molecules to mediate analogous principle for self-recognition and self–non-self discrimination during neuronal arborization. However, in contrast to fly Dscam1 isoforms which was shown to act as *cis*-monomer, sDscam and Pcdh proteins act as *cis*-multimeric recognition units to expand adhesive interfaces.

By contrast, sDscam-mediated self-recognition is analogous to that of Pcdhs, which is mediated by a mechanism coupling *cis* and *trans* interactions ^22,23,24,25, 29^. In vertebrate Pcdhs, ectodomains recognize each other via variable EC1/EC4 and EC2/EC3 *trans* homophilic interactions ^23,24,25,26,27,28,29^. Compared with the four EC domain-mediated *trans* interfaces in Pcdhs, the Chelicerata sDscam recognition interface contains only one Ig1 domain (or possibly with Ig2). Considering the fact that sDscams have a smaller proportion of *trans* interfaces, albeit with a larger *cis* interface, than Pcdhs, we speculate that stronger *cis* multimerization of sDscams compensates for their smaller *trans* adhesive interface to facilitate formation of stable *cis*/*trans* assembly complexes.

### Proposed model of sDscam interactions in self–non-self discrimination

Based on a large body of experimental evidence and structural modeling, we propose that sDscams form promiscuous *cis*-multimers at the cell surface that associate specifically in *trans* via an independent Ig1 self-binding interface. Similar to mouse Pcdhs ^24, 25, 27,28,29^, it is likely that full-length sDscam ectodomains in solution form a discrete array of multimers through specific *trans* dimerization of *cis* multimeric recognition units. However, as in Pcdhs ^23, 29^, such a structure cannot explain neuronal self–non-self discrimination because neurons with many neurites encounter insufficient diversity for self-recognition. Obviously, this self-recognition problem would be more severe for Chelicerata sDscams, which formed higher-order multimers than Pcdhs. Therefore, we speculate that Chelicerata sDscams could not adopt a discrete *trans* dimer of a *cis* multimeric assembly, but a zipper-like structure coupling *cis* and *trans* interactions like mouse Pcdhs ^24,25,26,27,28,29^.

However, in contrast to Pcdh *cis*-dimeric recognition units ^24,25,26,27,28,29^, sDscams can form *cis* tetramers and higher order multimers (Fig. 7d). Therefore, using sDscam *cis* tetramers as an example, we propose a two-dimensional latticed assembly structure model to account for sDscam-mediated cell-cell recognition (Fig. 7e). In this model, each sDscam *cis* tetramer could interact with multiple *cis* tetramers on apposed cell surfaces via independent *trans* Ig1/Ig1 self-binding, thereby forming a connected latticed assembly of proteins between cells. This model seems to be a “hybrid” structural framework between fly Dscams and vertebrate Pcdhs for Chelicerata sDscams, in which the Ig1/Ig1 self-binding *trans* interface is analogous to that of Ig7 of fly Dscams, while sDscam membrane-proximal *cis*-multimeric interfaces tended to largely resemble to that of vertebrate Pcdhs.

This model provides a reasonable explanation for how neurites discriminate self from non-self in Chelicerata, although detailed structures are not available. The presence of identical sDscam isoforms in two neurites of the same neuron would enable individual Ig1s of *cis*-multimers on apposed cell surfaces to self-bind each other independently of the composition of *cis*-multimers. Thus, the *trans* dimerization of *cis*-multimers could lead to a dense and connected lattice assembly between two apposing cell-surfaces, triggering strong homophilic interactions and inducing neurite repulsion (Fig. 7f). By contrast, because two neighboring neurons are incapable of expressing the same set of sDscam isoforms, mismatched Ig1s would lead to a scattered or sparse connected lattice assembly between apposing cell surfaces, triggering weak homophilic interactions. Thus, the resulting downstream signaling is below the threshold level and fails to initiate neurite repulsion (Fig. 7f). This model provides an evolutionary rationale for the smaller isoform number in Chelicerata by at least two orders of magnitude than that in flies. Although there are only ∼ 100 distinct sDscam isoforms in Chelicerata ^32^, this vast repertoire of combinatorial recognition specificities is sufficient to provide each neuron with a unique identity to discriminate between self and non-self. The major challenge for future studies will be to develop genetic techniques for Chelicerata species through which the sDscam diversity in the nervous system can be artificially manipulated.

### Chelicerata sDscams show more parallels with vertebrate Pcdhs than *Drosophila* Dscam1

Our findings indicate that Chelicerata *sDscam*s have striking parallels with *Drosophila Dscam1* and vertebrate *Pcdh*s, suggesting analogous roles (Fig. 8). Three encode large numbers of neuronal transmembrane protein isoforms; the individual isoforms are expressed stochastically and combinatorially, and the encoded proteins interact homophilically (Fig. 8) ^2, 3, 14^. In addition, such striking isoform diversity appears to underlie neuronal self–non-self discrimination, at least for the well-characterized *Drosophila* Dscam1 and vertebrate Pcdhs ^11, 12, 30, 31, 45^. Our results support and extend the notion that different phyla used different molecules or mechanisms to underlie analogous principle for mediating self-recognition and self-avoidance during neuronal arborization (Fig. 8) ^1, 2^. It would be interesting to determine which molecules mediate self-avoidance in other invertebrate phyla in the evolutionary gap between arthropods and vertebrates, particularly those lacking extensive Pcdh or Dscam diversity (e.g., the lancelet *Branchiostoma floridae*).

From an evolutionary viewpoint, Chelicerata sDscams are closely related with *Drosophila* Dscam1, but are not related with vertebrate Pcdhs ^3, 33^. However, in many respects, Chelicerata sDscams have more parallels with vertebrate Pcdhs (Fig. 8). Both are organized in a tandem array in the 5’ variable region, encoding the same order of magnitude of isoforms (50∼100) via alternative promoters. Both have a similar structural composition comprising six extracellular domains, a single transmembrane domain, and a cytoplasmic region. We have now shown that scorpion sDscam, like mouse Pcdhs, exhibited combinatorial recognition specificities based on the assembly of *cis-*multimeric recognition units, thereby sharing similar neuronal self-recognition logic with vertebrate Pcdhs. Thus, our findings further blur the distinction between the self-avoidance of invertebrates and vertebrates. It will be interesting to learn if convergent examples for self-avoidance in other animals are available. Finally, based on the remarkable parallels between Chelicerata sDscams and vertebrate Pcdhs, we wonder whether cadherins, in the animal kingdom, generate extraordinary isoform diversity via alternative splicing like their fly Dscam1 counterparts. One thing is certain—insight from extraordinary isoform diversity continues to deepen our understanding of basic biological principles.

## Methods

### Cell lines

Sf9 cells (a gift from Jian Chen, Zhejiang Sci-Tech University) were cultured in Sf-900™ II SFM (GIBCO, 10902088) supplemented with 10% fetal bovine serum (GIBCO, 10099141), and 1% Penicillin-Streptomycin (GIBCO, 15140163) at 27℃.

### Plasmid construction

DNA fragments encoding full-length sDscam isoforms or isoforms lacking the cytoplasmic domain were amplified by PCR using cDNA isolated from the scorpion *Mesobuthus martensii*^33^. PCR products were cloned into the pEasy-blunt zero cloning vector (TransGen Biotech, CB501-01) and ligated into the pFastBacHTB-mCherry/EGFP expression vector with appropriate restriction enzyme sites to guarantee that the opening reading frame was correct and the tags of EGFP and mCherry were fused to the C terminal of target proteins. To generate pFastBacHTB-mCherry and pFastBacHTB-EGFP vectors, the full-length mCherry amplified from the pmCherry-N1 vector (a gift from Xinhua Feng, Zhejiang University) and EGFP amplified from the pEGFP-N1 vector (a gift from Naiming Zhou, Zhejiang University) were inserted into the pFastBacHTB vector (a gift from Xiaofeng Wu, Zhejiang University) using the KpnI (forward primer) and HindIII (reverse primer) restriction sites. Domain deletion and substitution recombinant pFastBacHTB-mCherry vectors were made by using overlapping PCR. Extracellular-Myc-tagged sDscam vectors inserting the c-Myc (EQKLISEEDL) tag after the FNIII3 domain of sDscam were also generated by using overlapping PCR. The single mutation, double mutations and the mutations between close sDscam pairs were generated by site directed mutagenesis (Quikchange method). To obtain the pFastBac1-Myc/HA vector, sequences encoding the Myc (EQKLISEEDL) / HA (YPYDVPDYA) peptide were synthesized and annealed to form duple strand and then cloned into the pFastBac1 vector (a gift from Chuanxi Zhang, Zhejiang University) using restriction sites SphI and KpnI. pFastBac1-Myc truncations were obtained by using overlapping PCR. All recombinant vectors were confirmed by DNA sequencing. Ig and FNIII domain were predicted using the PROSITE (https://prosite.expasy.org/). Signal peptides (SP) and transmembrane domains (TM) were predicted using SMART (http://smart.embl-heidelberg.de/). Primer sequences used for PCR amplifications will be provided upon request.

### Antibody generation

Mouse monoclonal antibody against *M. martensii* sDscamβ6V2 (amino acids M1-L198) was generated by the Huabio. β6V2 antigen was cloned into pET-28a and transformed into Rosetta (DE3) *E. coli*, and purified using Ni-NTA beads (Smart-lifesciences) according to standard protocol.

### Recombinant baculovirus production

Baculoviruses were obtained according to the manufacturer’s instructions of Bac-to-Bac Baculovirus Expression System (GIBCO, 10359016). Briefly, to generate a recombinant bacmid, the pFastBac plasmid was transformed into DH10Bac competent cells (Biomed, BC112) and blue-white screening was used to pick the white colonies and recombinant bacmid DNA was analyzed by PCR. Recombinant bacmid was transfected into Sf9 cells using Lipofectamine 3000 Reagent (Invitrogen, L3000015). Cells were incubated at 27°C until see signs of viral infection. Then we harvested the virus from the cell culture medium to get P1 baculovirus. P1 viruses were added to Sf9 cells grown in the 6-well plates, after 72 h of incubation at 27°C, the cells were centrifuged to obtain the supernatant as P2 viruses. All baculoviruses should be stored at 4°C.

### Cell aggregation assays

Cell aggregation assay was performed as previously reported with little modification ^34^. Sf9 cells grown in each well in 6-well plates were infected with P2 viral of target proteins tagged with mCherry or EGFP 10∼30 ul and incubated at 27°C for 3 days. Cells were suspended and transferred to the 2ml tube, centrifuged at 1,000rpm for 5 min. The supernatants were discarded, and the cell pellets were resuspended with the 1ml 1×HCMF (1:10, Leagene Biotechnology, CC0073) gently and centrifuged at 1,000rpm for 5 min again. Cells were then resuspended with 1ml 1×HCMF gently. For cell aggregation assay, 400ul cell suspension was transferred into each well in 6-well plates containing 2ml 1×HCMF per well. For binding specificity assay, 200ul cell suspension of each sample was transferred into each well. The 6-well plates used in the cell aggregation assay firstly was added the 1%BSA in 1×HBSS (1:10, Gibco, 14185052) at 4°C overnight, then washed once with D-PBS and 2ml 1×HCMF was added to each well. Cell suspension in 6-well plates was incubated at 27°C with gyratory shaker (IKA KS260) at 60rpm for 30 min. Finally, images were captured using the Nikon Ti-S inverted fluorescence microscope.

### Binding specificity assay for cells expressing single or multiple sDscam isoform(s)

Differentially tagged sDscam isoforms were infected into Sf9 cells as described above. We observed that sDscam’s surface expression is different among sDscamα or sDscamβ isoforms in the coexpression experiments. Thus, sDscamα-sDscamα or sDscamα-sDscamβ were used in appropriate ratio roughly guaranteed the approximate equal surface expression. Images were captured using the Nikon Ti-S inverted fluorescence microscope, and the aggregates containing red cells only, green cells only, and both red and green cells (Red-Green) were observed and counted for analysis of binding specificity.

### Quantification of the size of cell aggregates using matlab

The images of cells from three independent aggregation experiments were used to quantify the relative size of cell aggregates generated by each sDscam isoform. The images were first converted to black and white formats with 2160ⅹ2560 pixels. Objects with 1000 or fewer pixels were categorized as small (<10 cells), objects between 1000 pixels and 6000 pixels were categorized as medium (10 to 80 cells), and objects larger than 6000 pixels were categorized as large (>80 cells). The number of aggregates of each size category was then counted for analysis.

### Immunostaining

Sf9 cells were seeded onto the coverslips (WHB Scientific) coated with 1mM poly-L-lysine (Sigma, P6282) in 6-well plates, and were infected by the viruses from the P2 stocks of sDscamα14Δcyto, sDscamβ4v3Δcyto and sDscamα14ΔcytoΔIg1, these sDscam proteins inserted c-Myc tag between the FNIII3 and TM domain as well as carried a mCherry tag at the C terminus. after 72hr, cells were fixed with 4% Paraformaldehyde Fix Solution (PFA, Sangon Biotech, E672002-0500) for 20min at room temperature, then these non-permeabilized cells were washed three times with D-PBS (Sangon Biotech, E607009). Cells were blocked with 5% BSA in PBS and incubated with anti-Myc tag monoclonal antibody (1:400, Earthox, E022050-01) overnight at 4 °C. Subsequently, cells were washed three times with PBS, incubated with goat Anti-Mouse IgG (H+L) Dylight488 (1:500, Earthox, E032210-01) diluted in 5% BSA of D-PBS for 1∼2 h at room temperature and then washed three times with D-PBS. Finally, cell was stained by Hoechst (2μg/ml, invitrogen, Hoechst 33342) for 15∼30 min for nucleus staining and imaging with a laser scanning confocal microscope LSM800 (Carl Zeiss).

### Co-immunoprecipitation and western blot analysis

Sf9 cells grown in 6-well plates were infected with viral and incubated for 3 days at 27°C. Cells were lysed in the IP lysis buffer (Thermo Fisher Scientific, 87787). The cells were incubated in lysis buffer for 30 min at 4°C. The supernatant was collected by centrifugation at 13,000 g for 20 min at 4°C. Subsequently, cellular extracts were incubated with appropriate antibodies at 4°C overnight, followed by incubating with pre-washed protein A/G magnetic beads (Thermo Fisher Scientific, 88802) for 3 h at 4°C. The beads were collected and washed, then boiled with SDS sample buffer (Sangon Biotech, C508320-0001) for 10 min. The beads were separated and then the supernatant was saved for western blotting according to the standard methods. In brief, proteins were electrophoresed in Tris-Glycine gel (Sangon Biotech, C651101-0001) and transferred to the PVDF membranes (Millipore, IPVH00010). Membranes were incubated with antibodies as described and then performed protein signal detection. For multimer detection of sDscam, target proteins in Sf9 cell and tissues were extracted using the RIPA lysis buffer (strong) (Cowin Biosciences, CW2333S) freshly supplemented with 1% protease inhibitor cocktail. The sample was treated with nonreducing sample buffer (Sangon Biotech,C516031) without boiling then directly centrifuged at 13,000rpm for 20 min at 4°C. The proteins were electrophoresed in Tris-Glycine PAGE gel (Sangon Biotech, C651104-0001). Then the western blotting was performed as described above. The anti-Mma β6V2 monoclonal antibody (produced by Huabio, Hangzhou) raised against the Ig1 domain of Mma β6V2 was used by tissue western blotting.

The antibodies used in IP: anti-HA Tag rabbit polyclonal antibody (1:50, Earthox, E022180-01), HA-Tag (C29F4) rabbit mAb (1:50, Cell Signaling Technology, 3724S), anti-GFP rabbit polyclonal antibody (1:50, Earthox, E022200-01). The primary antibodies used in Western blotting: anti-HA Tag monoclonal antibody (1:5000, Earthox, E022010-01), anti-Myc Tag antibody (1:5000, Earthox, E022050-01), anti-mCherry Tag antibody (1:5000, Earthox, E022110-01), anti-GFP Tag mouse monoclonal antibody (1:5000, Earthox, E022030-01), Anti-sDscamβ6V2 Monoclonal Antibody (1:1000). The secondary antibodies used in Western blotting: HRP AffiniPure goat anti-Mouse IgG (1:8000, Earthox, E030110-01).

### Homology modeling and protein-protein docking

Ig1 homodimeric models of sDscam were built with swiss-model (https://www.swissmodel) using Dscam Ig7 as a temple ^39, 40, 46^. Macromolecular interface was furthermore explored using PDBePISA (https://www.ebi.ac.uk/msd-srv/prot_int/cgi-bin/piserver) ^47^, the results show some buried residues and their potential interaction (hydrogen/disulphide bond, salt bridge or covalent link), single and double complementary mutations of these candidate residues were conducted upon this. Specificity-determining residues of six sDscamα pairs were screened also based on homology modeling. Similarly, TM domain of sDscamβ6v2 was generated by swiss-model, and all these structure figures were prepared with VMD package (https://www.ks.uiuc.edu/Research/vmd/). In addition, Ig3-FNIII3 domain of sDscamβ6v2 was predicted as monomer by swiss-model and then was used for homologous dimer docking by ZDOCK server (http://zdock.umassmed.edu/m-zdock/). Finally, the dimer interface were visualized with the PyMOL package (www.pymol.com).

### Sequence alignments and heatmap analysis

Multi-sequence alignments of Ig1 and Ig1-2 domains of the sDscam were carried out with Clustal Omega (https://www.ebi.ac.uk/Tools/msa/clustalo/) and Align (https://www.uniprot.org/align/). Then the sequence similarity heatmap was made by R language.

### Statistics

Statistical significance was calculated by using IBM SPASS Statistics V22.0 (Mann-Whitney U-test) to determine significant difference of cell aggregation size between TM domains shuffled isoforms, as well as β6v2FL-Cherry and β6v2Δcyto-Cherry isoforms. The number of independent experiment of duplicates, the statistical significance, and the statistical test are indicated in each figure or figure legend where quantification is reported.

## Author contributions

YJ conceived of this project. FZ, GC, CD, GL, HL, and BX designed and performed the experiments; FZ, GC, CD, GL, and HL conducted cell aggregation assays and binding specificity assay; FZ, GC, CD and GL performed multimerization assay; GC conducted sDscam expression and antibody generation; GC, CD, GL, and FZ, performed co-immunoprecipitation and western blot analysis; FZ, SH and ZD performed homology modeling and protein-protein docking; YJ, FZ, BX, FS, YD and XY analyzed the data; YJ and FZ wrote the manuscript; all authors discussed the results and commented on the manuscript.

## Acknowledgements

This work was supported by research grants from the National Natural Science Foundation of China (31630089, 31430050, 91740104).

