## supplemental information for "Dscam homophilic specificity is generated by high order *cis*-multimers coupled with *trans* self-binding of variable Ig1 in Chelicerata"

**This PDF file includes:**

Supplementary Fig.1 to 7

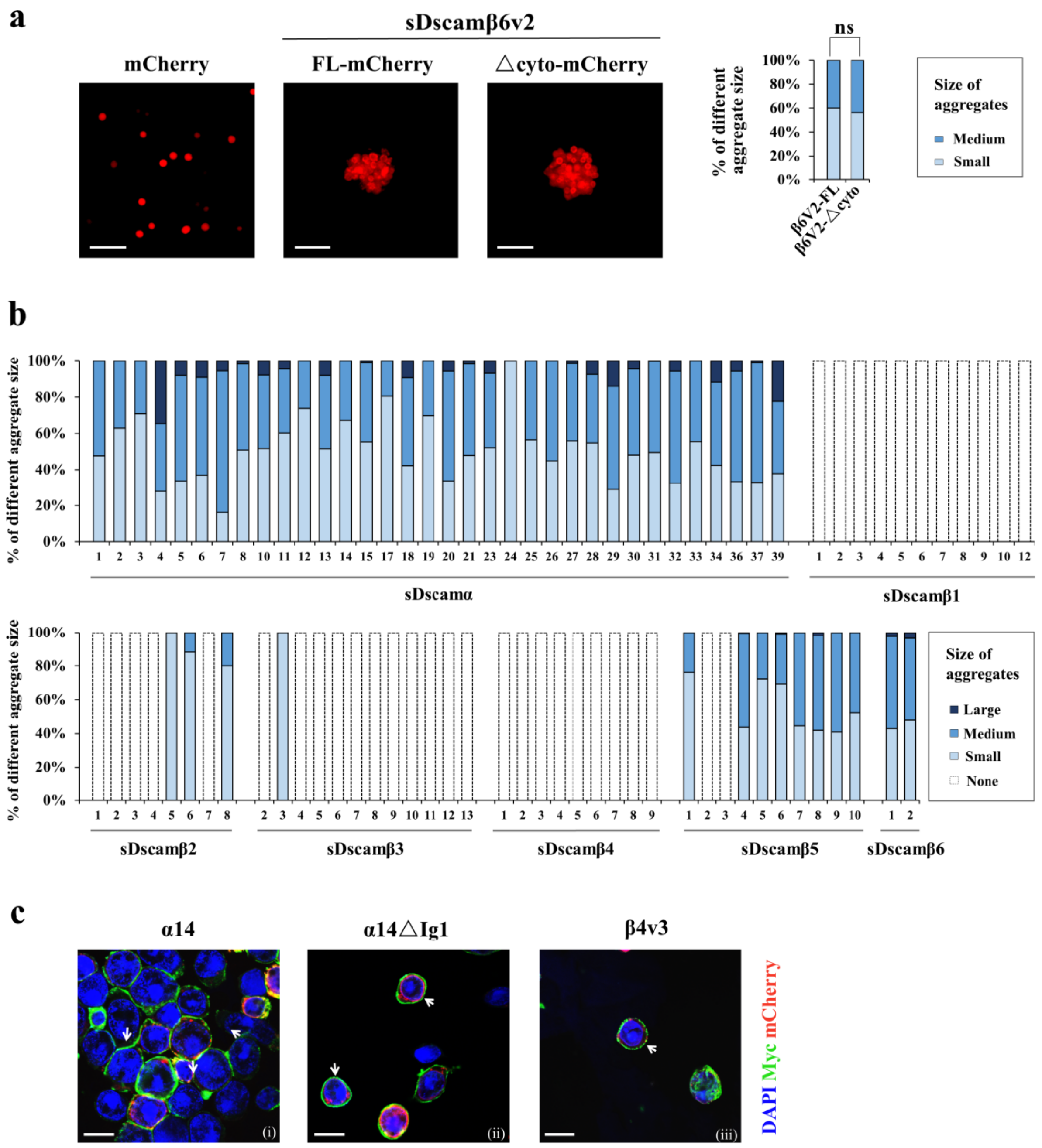


**Supplementary Fig. 1 Quantification of the size of sDscam-mediated cell aggregates. Related to Fig. 1.**

(**a**) Cell aggregation mediated by sDscam isoforms. Sf9 cells expressing Cherry, full-length sDscamβ6v2 (β6v2FL-Cherry), or sDscamβ6v2 lacking the cytoplasmic region (β6v2Δcyto-Cherry) were prepared for cell aggregation assay. This result showed that deletion of the cytoplasmic region of sDscam had no detectable effect on the formation of cell aggregates.

Scale bar, 100μm.

(**b**) Quantification of the sizes of cell aggregates. A bar graph showed the size distribution of cell aggregates with cells expressing individual sDscam isoform. Cells expressing each of alternative sDscamα and sDscamβ5–β6 isoforms showed extensive aggregation in all isoforms tested. By contrast, sDscamβ1–β4 isoforms failed to mediate cell aggregation, except for a few isoforms. The cell aggregates were classified according to the number of cells they contain — small size (<10 cells), medium size (10–80 cells), and large size (>80 cells). The results of cell aggregation were obtained from three independent experiments.

(**c**) Cell surface expression of sDscam. Sf9 cells expressing mCherry-tagged sDscamα14 (left panel), sDscamβ4v3 (right panel) and sDscamα14△Ig1 (medium panel) constructs fused an extracellular Myc tag were assayed for cell surface expression. White arrows indicate Myc staining at cell-cell contacts. These data indicate that sDscamα, sDscamβ and sDscamα14△Ig1 proteins could be localized on the cell surface. Scale bar, 20μm.

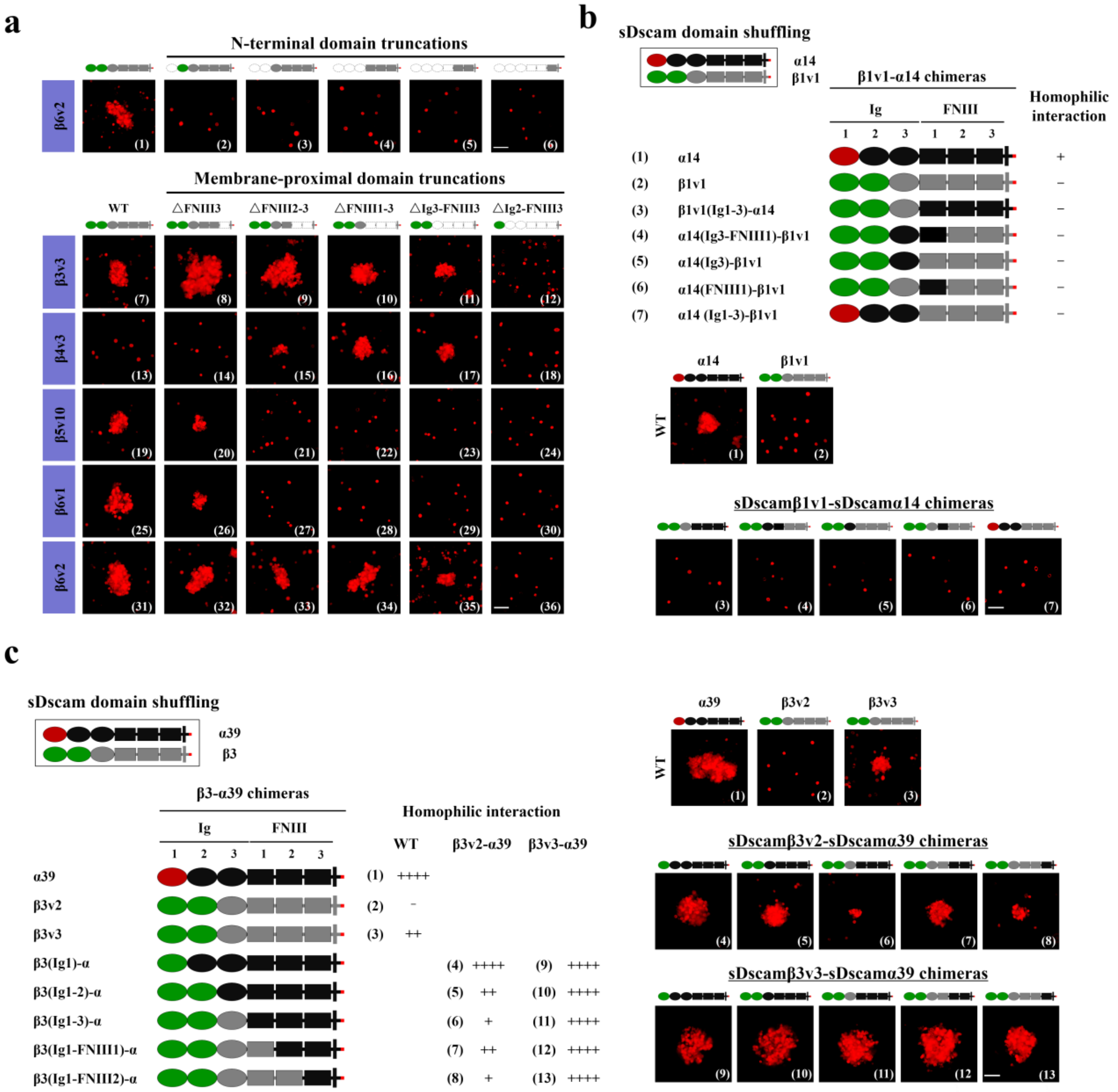


**Supplementary Fig. 2 Homophilic *trans*-binding was regulated by variable and constant domains of sDscam. Related to Fig. 2.**

(**a**) A series of N-terminal (upper) and membrane-proximal (lower) truncations of the extracellular domains of sDscam fused with mCherry were examined for cell aggregation assay. These results indicate that the first two N-terminal domains are required for *trans* homophilic binding.

(**b**) Schematic diagrams of domain shuffling between sDscamα14 and sDscamβ1v1, along with the results of homophilic binding. Extracellular domains of sDscamα14 (black) were replaced with the corresponding domains of sDscamβ1v1 (gray), or vice versa. These results indicate that either individual or combined replacement of constant extracellular domains of sDscamβ1v1 by corresponding region of sDscamα14 did not cause cell aggregate (panels 3–7).

(**c**) Homophilic binding assays for domain shuffling between sDscamα39 and sDscamβ3v2/v3. Schematic diagram along with a summary of aggregation outcomes was shown on the left panel, while cell adhesion images corresponded to the shuffled chimeras were showed on the right panel. Extracellular domains of sDscamα39 (black) were replaced with the corresponding domains of sDscamβ3 (gray), or vice versa.

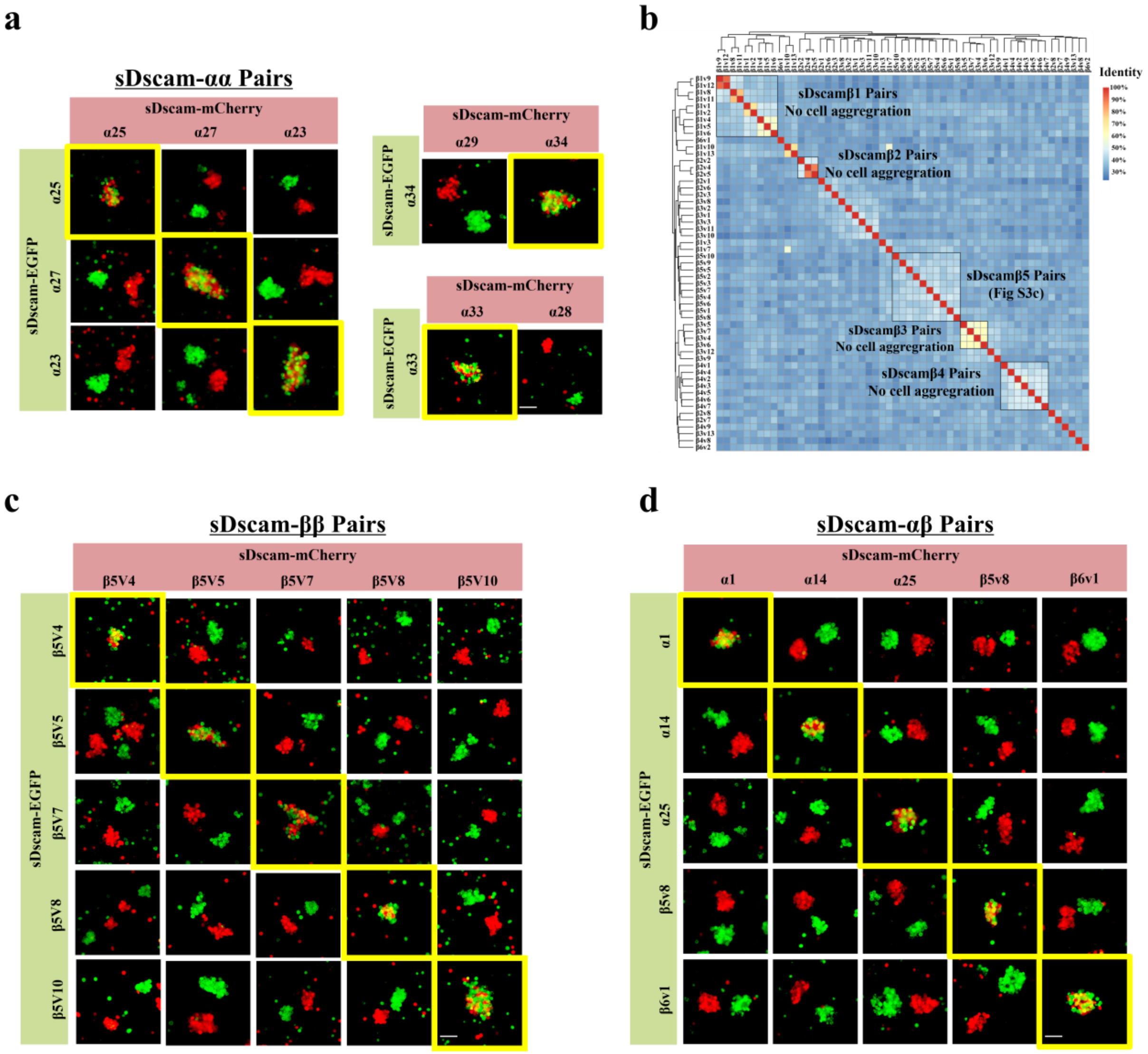


**Supplementary Fig. 3 sDscam isoforms engaged in highly specific homophilic interactions. Related to Fig. 3.**

(**a**) Pairwise combinations within representative sDscamα were assayed for their binding specificity. Subsets of these isoforms were marked within the boxed region in Fig. 3b. Scale bar, 100 μm.

(**b**) Heat map of pairwise protein sequence identities of the Ig1 domains of sDscamβ isoforms and their evolutionary relationship were presented. Subsets of the isoforms within the boxed region were assayed (sDscamβ5 subset).

(**c, d**) Pairwise combinations within representative sDscamβ&β and sDscamα&β were assayed for their binding specificity. Scale bar, 100 μm.

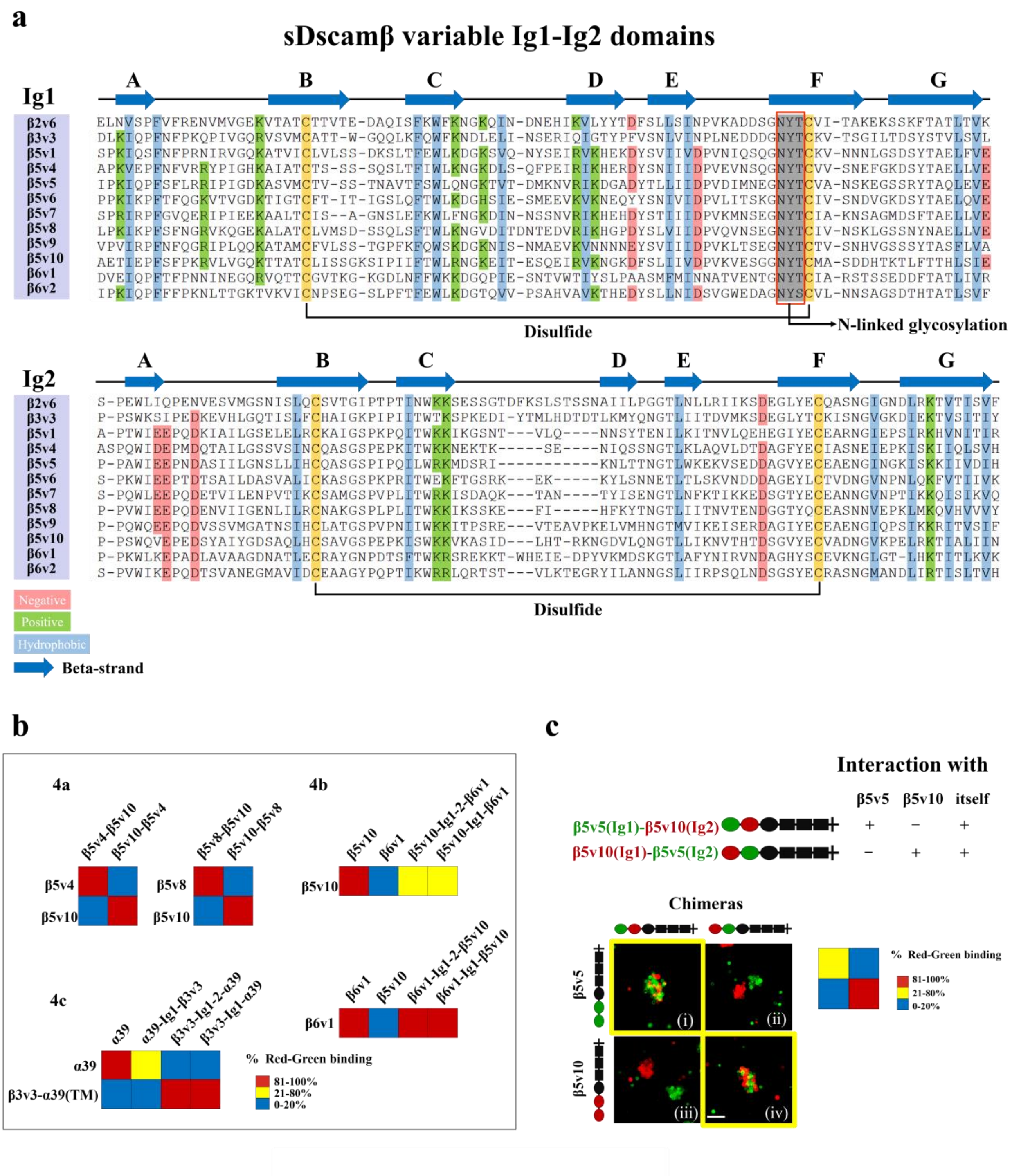


**Supplementary Fig. 4 sDscam *trans*-binding specificity is mediated mainly by Ig1 domain. Related to Fig. 4.**

(**a**) Amino acid sequence alignment of variable Ig1-Ig2 domains of sDscamβ isoforms. The locations of β strands are indicated above the alignment, and the conserved hydrophobic residues were highlighted in blue, the conserved negative residues were highlighted in red, as well as the conserved positive residues were highlighted in green. Potential disulfides within Ig domains are indicated. The predicted N-linked glycosylation motifs are shaded in gray and within the red box.

(**b**) Coaggregation results of two labeled cell populations. The percentages of red and green cell coaggregates from Fig. 4a–c were illustrated as a heat map.

(**c**) Domain-shuffled chimeras of sDscamβ5 isoforms and their parental counterparts were assayed for binding specificity. The results indicate that the presence of a single common Ig1 domain is sufficient to confer co-aggregation between sDscam isoforms.

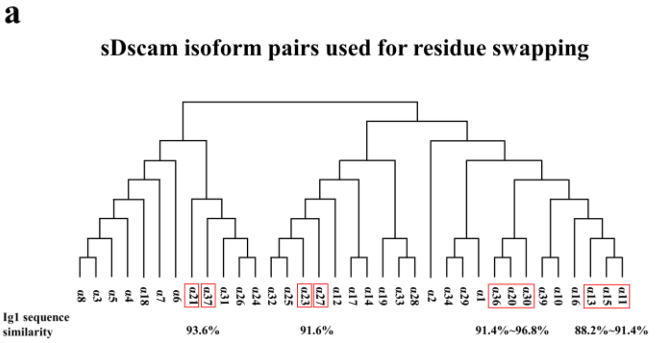


**b**

**Residues swapped table**

| sDscam isoforms | Residues swapped | Residue identities | Image |
|---|---|---|---|
| α30&α36&20 | 1, 15, 21, 22 | α30: S1, S15, V21, T22<br>α36&α20: P1, N15, I21, I22 | Fig 5c |
| α21&α37 | 5, 10, 56 | α21: R5, T10, E56<br>α37: K5, A10, D56 | Fig 5d<br>Fig S5g |
| α11&α15 | 22 | α11: I22<br>α15: T22 | Fig S5c |
| α13&α15 | 22 | α13: I22<br>α15: T22 | Fig S5c |
| α11&α13 | 26, 52 | α11: N26, V52<br>α13: I26, M52 | Fig S5d<br>Fig S5e |
| α23&α27 | 6, 19, 52 | α23: Q6, Q19, L52<br>α27: P6, E19, V52 | Fig S5e<br>Fig S5f |

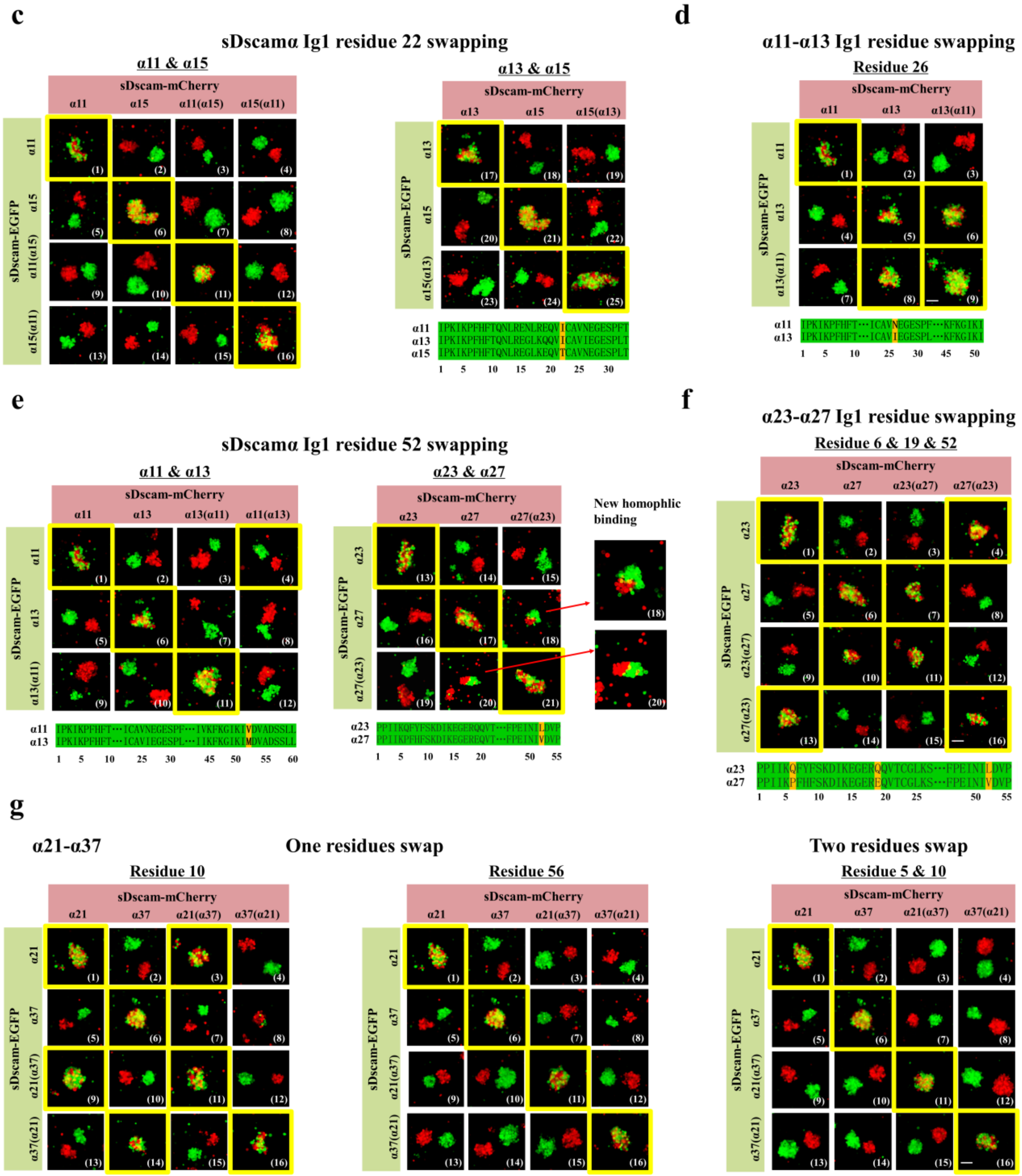

**Supplementary Fig. 5 Identification of Ig1 specificity-determining residues. Related to Fig. 5.**

(**a**) sDscam isoform pairs used for residue swapping. A phylogenetic tree depicted the relationships of Ig1 domains, whereas pairs of Ig1 domains for residue swapping were marked by red rectangle.

(**b**) The table outlines the swapped residues between sDscam isoform pairs.

(**c**) Swapping single residue I22T of Ig1 between sDscamα11 and α15, or between sDscamα13 and α15 was not sufficient to swap specificity, but produced new specificity.

(**d**) Swapping single residue N26I of Ig1 between sDscamα11 and α13 had no effect on specificity.

(**e**) Swapping single residue M52V of Ig1 between sDscamα11 and α13 was not sufficient to swap specificity, but produced new specificity.

(**f**) Three residues 6, 19 and 52 swapping of Ig1 between sDscamα23 and α27 fully switched binding specificity.

(**g**) Cell aggregation assays of isoforms containing wild-type and other residue-swapped isoforms between sDscamα21 and α37, supplemental to Fig. 5d. Swapping of either one of three residues did not affect binding specificity (left penal) or produced new binding specificity (middle penal), while swapping of two of three residues produced new binding specificity (right penal).

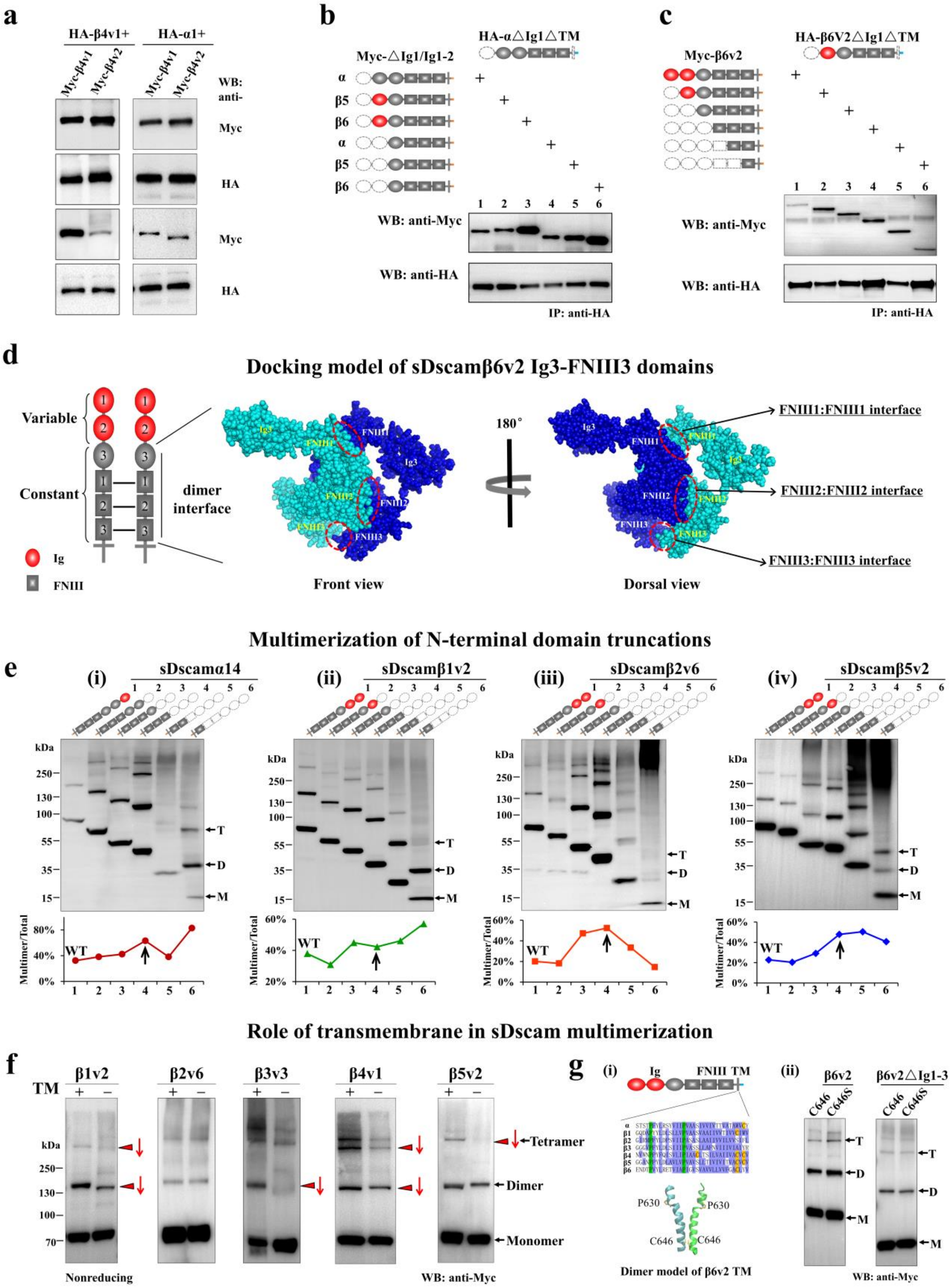


**Supplementary Fig. 6 sDscams isoforms form high order *cis*-multimers. Related to Fig. 6.**

(**a**) Co-IP assay between various sDscam isoforms. Myc-tagged sDscamβ4 proteins could strongly coimmunoprecipitate with HA-β4v1 and HA-α.

(**b**) sDscamαΔIg1ΔTM were immunoprecipitated strongly with a series of other sDscamΔIg1 or sDscamΔIg1–2 proteins.

(**c**) *Cis* interaction could occur in the absence of N-terminal domains. HA-β6v2ΔIg1ΔTM and different Myc-tagged sDscam isoforms were immunoprecipitated using anti-HA antibody and probed with anti-Myc or anti-HA antibodies. This result indicated that robust coimmunoprecipitation between HA-sDscamβ6v2 and Myc-sDscamβ6v2 should not result from *trans* interactions, but *cis* interactions.

(**d**) Docking model of sDscamβ6v2 constant domains. In this docking model, sDscam homodimer exhibited multiple parallel interfacial regions involving FNIII1–3 domains (red dashed circle).

(**e**) A series of N-terminal truncations of the extracellular domain were examined for multimerization assay in sDscamα14 (i), sDscamβ1v2 (ii), sDscamβ2v6 (iii) and sDscamβ5v2 (iv). Graphs of the ratio change of multimer/total were shown below.

(**f**) Deleting TM domain could greatly reduce the efficiency of multimerization. Red arrow marks changed multimers, and downward red arrow marks the decrease of multimerization efficiency.

(**g**) Effect of TM cysteine mutations on *cis*-multimerization. (i) Sequence alignment of sDscam TM domains was shown in left panel, whereas sDscamβ6v2 TM domain showed contiguous-helix formation, including Cys646. (ii) Cysteine residue mutation in TM domain did not affect the formation of multimers.

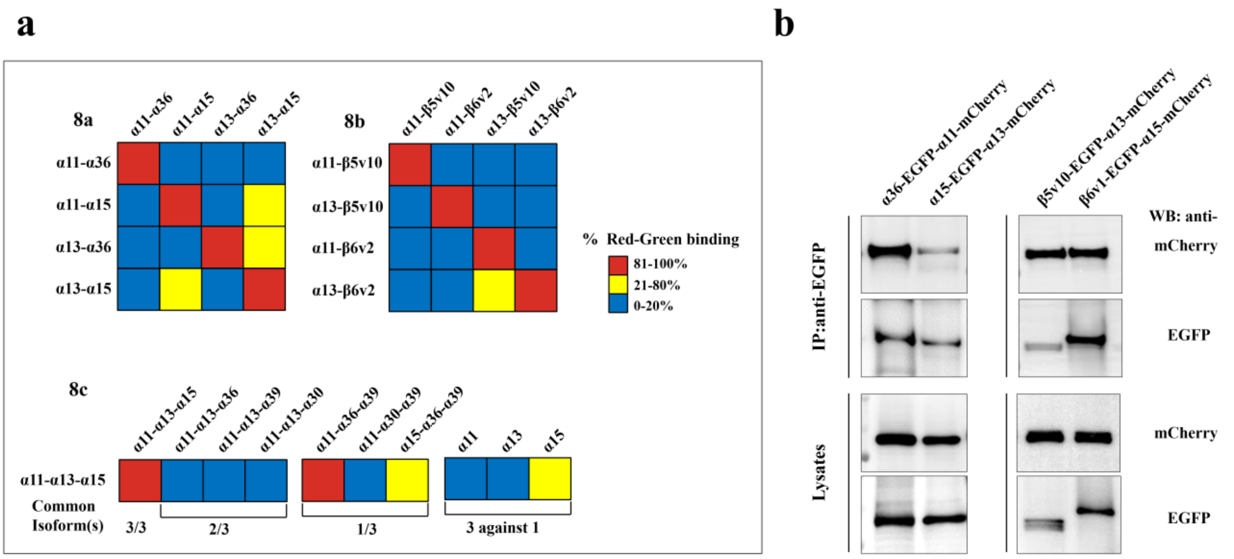


**Supplementary Fig. 7 Chelicerata sDscams show more parallels with vertebrate Pcdhs than *Drosophila* Dscam1. Related to Fig. 7.**

(**a**) Coaggregation results of two differentially labeled cell populations. The ratios of red and green coaggregates from are shown as a heat map.

(**b**) All sDscamα and sDscamβ isoforms tested (Fig. 7a–c) interacted strongly with each other in co-IP experiments. Lysates from Sf9 cells cotransfected with sDscam fused a C-terminal EGFP-tag and different mCherry-tagged sDscam isoforms were immunoprecipitated using anti-EGFP antibody and probed with anti- mCherry or anti- EGFP antibodies.
